# D3 nutraceutical pre-supplementation ameliorates behavioral, ionotropic receptors, and synaptic proteins alterations in a MK-801 induced mouse model of schizophrenia

**DOI:** 10.64898/2026.06.19.733409

**Authors:** Omalur Eshwari, Kishore Golla, Shashikant Patel, Vijay Aravind Yenagi, Roli Kushwaha, Sharon Mariam Abraham, Sumana Chakravarty, Raghavendra B. Nayak, Pragya Komal

## Abstract

Vitamin D3 (VD) deficiency is a global health concern, and its supplementation has been shown to alleviate inflammation and oxidative stress across numerous neurological disorders. However, the beneficial effect of this common nutraceutical in schizophrenia (SCZ) remains inadequately explored. The present study investigated the presupplementation effects of VD on positive and cognitive symptoms in a MK-801induced mouse model of SCZ. MK-801, a non-competitive NMDA receptor antagonist, is a widely used drug that mimics some of the psychotic symptoms associated with SCZ. The repeated administration of a single dose of MK-801 (0.5mg/kg; intraperitoneally) for two weeks produced hyperlocomotion, anxiety- like behavior, and working memory deficits in MK-801-induced SCZ-like mice. These behavioral abnormalities were significantly attenuated in VS5 mice (SCZ mice presupplemented with 500 IU/kg/day of VD). At the molecular level, VD rescued gene expression of major NMDA receptor subunits (NR1, NR2A, NR2B), α7 nicotinic acetylcholine receptors (α7nAChRs), and neurotrophin factors (NGF and BDNF). A restoration of PSD-95 protein expression, accompanied by downregulation of calcineurin, was also observed in the prefrontal cortex (PFC) of VS5 mice, suggesting protective effects of VD on synaptic communication and function in SCZ. In vitro studies showed that calcitriol (1 μM) treatment of HEK-293 T cells transfected with α7nAChRs potentiated the single-channel current amplitude and demonstrated a direct modulatory effect of this nutraceutical on α7nAChRs expression and function. In silico JASPAR analysis further identified putative Vitamin D response elements (VDREs) within the promoter regions of various target genes, supporting the genomic action of VD. Additionally, VD deficiency was observed in Indian SCZ patients, highlighting its potential clinical relevance. Together with our previous findings (Manjari et al., 2022, 2023), the present study also demonstrates anti-inflammatory, anti-cholinesterase, neurotrophic, and synaptic-enhancing effects of VD, deepening our understanding of the multifaceted neuroprotective effects of the “D3” neurosteroid in neuropsychiatric disorders such as SCZ.

**Highlights:**

- VD presupplementation improves the behavioral deficits in MK-801 induced SCZ mice.
- Nutraceutical intervention normalizes the gene expression of major NMDARs subunits namely, NR1, NR2A, NR2B, in the PFC of SCZ mice.
- VD mediates a restoration in the expression and function of α7nAChRs in SCZ mice.
- VD exhibits neuroprotective, neurotrophic, synaptoprotective, anti-inflammatory and anti-acetylcholinesterase effects, highlighting its therapeutic potential in SCZ.

## Background

An intricate relationship between nutraceutical interventions like Vitamin D3 (VD) in mental health disorders has recently been extensively acknowledged in the field of neuropsychiatric and neurological disorders [1–13]. VD deficiency (VDD) is a global pandemic and has been associated with adverse neurodevelopmental outcomes and predominant psychotic symptoms like those observed in schizophrenia (SCZ) [14–22]. Clinical studies have reported a strong association of VD deficiency with SCZ [16, 23]. SCZ is a neuropsychiatric disorder with multiple manifestations in which glutamate receptor hypofunction theory emerges from various neuroimaging methodologies, transcriptomics, proteomics studies, and postmortem brains from patients [24–30]. The psychosis symptoms observed in SCZ can also develop due to a combination of altered environmental factors, genetics, socioeconomic conditions, stress, neuroinflammation, maternal immune activation, use of stimulants, metabolic disorders and dysregulation of neurotransmitters in the various regions of the brain [31, 32]. Some central features underlying the disability in SCZ include synaptic dysfunction, deficits in cognition, and glutamatergic abnormalities [33–38]. The N-methyl-D-aspartate (NMDA) type of glutamate receptor antagonist, MK-801 (dizocilpine), is known to reproduce most of the symptomatic features of SCZ [39–41]. It is imperative to project that the compounds that can rescue dysfunctional glutamatergic neurotransmission, enhance or normalize NMDA receptor expression, and reduce cognitive deficits may represent a crucial central focus in the SCZ research field. In support of this inference, multiple studies have documented the therapeutic potential of VD in preclinical studies of neurodegenerative disorders [2, 3, 16, 42–51]. Vitamin D3 (VD) is a fat-soluble neurosteroid that has been postulated to have multifaceted functions in neurodegenerative disorders like Huntington’s disease (HD), schizophrenia (SCZ), Alzheimer’s disease (AD), Parkinson’s disease (PD), and Major depressive disorders (MDD) [3, 48, 50, 52–56]. A comprehensive understanding of simple nutraceutical interventions like VD on glutamatergic abnormalities, and psychosis symptoms induced by MK-801, however, remains limited.

In light of this aspect, we explored whether Vitamin D3 (VD, cholecalciferol) pre-administration could restore some of the neuropsychiatric symptoms induced by MK-801 (dizocilpine) in a mouse model of SCZ. VD is a neurosteroid and can be produced from sun exposure in the skin or ingested from the diet and is transported to the liver by binding to Vitamin D binding protein (VDBP) [57]. It is then hydroxylated by the enzyme cytochrome P450 family 2 subfamily R member 1 (CYP2R1) or cytochrome P450 family 27 subfamily A member 1 (CYP27A1) that produces biologically inactive, 25-hydroxylated form of VD (25(OH)D3) or calcidiol [3]. The second hydroxylation, catalysed by 1-alpha-hydroxylase (CYP27B1), converts 25(OH)D3 to the biologically active form of VD referred as 1,25 dihydroxy Vitamin D3 (1,25(OH)2D3) or calcitriol [58]. Although this process takes place primarily in the kidneys, neurons and glial cells also expresses CYP27B1 gene that permits the local synthesis of the active form of VD within the central nervous system (CNS) [59]. It is well documented that the local effect of increased calciferols (calcidiol and calcitriol) concentration can bring myriad genomic and non-genomic effects of VD in the brain by binding to PDIA3 (also known as Protein Disulfide Isomerase Family A Member 3 or MARRS) or Vitamin D receptors (VDRs). These membrane and nuclear receptors are mainly responsible for the majority of VD’s physiological functions [59–62].

VD plays an essential role in neuroplasticity, neuroprotection, neurotransmitter biosynthesis, and neurotransmission in developing and adult brains [2, 52, 55, 63–66]. It is also known to reduce oxidative stress and inflammation caused by free radicals and reactive oxygen metabolites by upregulation in the gene expression of many neurotrophic factors [45, 67–72]. Previously, we confirmed the neuroprotective effects of VD in a 3-nitropropionic acid-induced mouse model of HD [45]. In the present study, we have taken 500IU/kg/day dose of VD, as this dose showed the maximum efficacy in alleviating the behavior symptoms as observed previously under neurotoxic conditions. Other doses of VD (1000 IU/kg and 2000 IU/kg) showed a similar effect on behavioral abnormalities induced by MK-801, with no significant difference from the 500IU/kg VD dose. Therefore, we focused primarily on elucidating the potential benefits of pre-supplementation effects of 500IU/kg VD dose in SCZ that were evaluated through various standard behavioral, biochemical, gene expression, and electrophysiological analyses. MK-801, the pharmacological antagonist for N-methyl-D-aspartate receptor, is a known inducer of ionotropic receptors alterations and some synaptic protein abnormalities in the prefrontal cortex^69–71^. In the present work, we found that the pre-supplementation intervention with a dose of 500IU/kg of VD decreased psychotic symptoms of SCZ, rescued mRNA expression of NMDA receptor subunits (NR1, NR2A and NR2B), α7nAChRs, inflammatory cytokines like tumor necrosis factor-alpha (TNF-α), neurotrophic factors (NGF, and BDNF), protein expression of Vitamin D receptor (VDR), and synaptic markers like PSD-95 in the prefrontal cortex of SCZ mice. Furthermore, *in-silico* JASPAR analysis identified putative VDRE’s in the promoter region of target genes, suggesting a potential VDR-mediated transcriptional mechanism. Our findings reinforce and extend the notion on neurotrophic, channeloprotectant, and synaptotrophic effects of VD in the area of mental health disorders [43, 73–75].

## Methods

### Animal Procurement

Eight to ten-week-old C57BL/6 male mice (average weight; 22 ± 3g) were obtained from ICMR-National Animal Resource Facility for Biomedical Research (ICMR-NARFBR), Hyderabad. The mice were placed in quarantine conditions at the Central Animal Facility, BITS- Hyderabad for 5-7 days. The mice were then housed in groups of 2-3 mice per cage on a 12/12-hour light/dark cycle at a constant temperature (24°C) and had free and continuous access to food and water. Frequent bedding change was performed, and feces were discarded regularly to prevent infections. All behavioral studies were conducted in the light phase of the cycle. Animal weights were measured daily throughout the experiment cycle to assess animal health (Fig. S1). All the animal experiments were carried out with the approval of the Institutional Animal Ethics Committee (IAEC), BITS – Pilani, Hyderabad, India (Protocol Approval Number: BITS-Hyd-IAEC-2022-34). All efforts were made to minimize animal suffering and to reduce the animal number used in this study.

### Experimental design

After an acclimatization period (quarantine) of 5-7 days, our experimental mice were randomly divided into different experimental groups as shown below:

**(i) Control**: Mice were administered an intraperitoneal (i.p) injection of 1X saline.
**(ii) SCZ**: MK- 801 (0.5 mg/kg/day) intraperitoneal (i.p) injections were given daily for 15 days [72] that induced SCZ-like symptoms.
**(iii) VD5; VD1 and VD2**: An intraperitoneal (i.p) injection of three different doses of only Vitamin D3 (VD) or cholecalciferol was given for the first two weeks (−1 day to −15 days, Fig.1).

a. **VD5**: Mice received only **500IU/kg/day** (or 12.5μg/kg/day) of VD for the first two weeks (−1 day to −15 days).
b. **VD1**: Mice received only **1000IU/kg/day of VD** for the first two weeks (−1 day to −15 days).
c. **VD2**: Mice received only **2000IU/kg/day of VD** for the first two weeks (−1 day to −15 days).
**(iv) VS5; VS1; VS2**: An intraperitoneal (i.p) injection of MK-801 (0.5 mg/kg/day) was given to mice presupplemented with three different doses of Vitamin D3 (VD) or cholecalciferol VD (500/ 1000/ 2000 IU/kg/day); as highlighted below:
**A. VS5:** Mice presupplemented with **500IU/kg/day of VD** for the first two weeks (−1 day to −15 days) were administered with an intraperitoneal (i.p) injection of MK-801 (0.5 mg/kg/day) for an additional two consecutive weeks (0^th^ day to 15^th^ day).
**B. VS1:** Mice presupplemented with **1000IU/kg/day of VD** for the first two weeks (−1 day to −15 days) were administered with an intraperitoneal (i.p) injection of MK-801 (0.5 mg/kg/day) for an additional two consecutive weeks (0^th^ day to 15^th^ day).
**C. VS2:** Mice presupplemented with **2000IU/kg/day of VD** for the first two weeks (−1 day to 15 days) were administered with an intraperitoneal (i.p) injection of MK-801 (0.5 mg/kg/day) for an additional two consecutive weeks (0^th^ day to 15^th^ day).

**Fig. 1.**
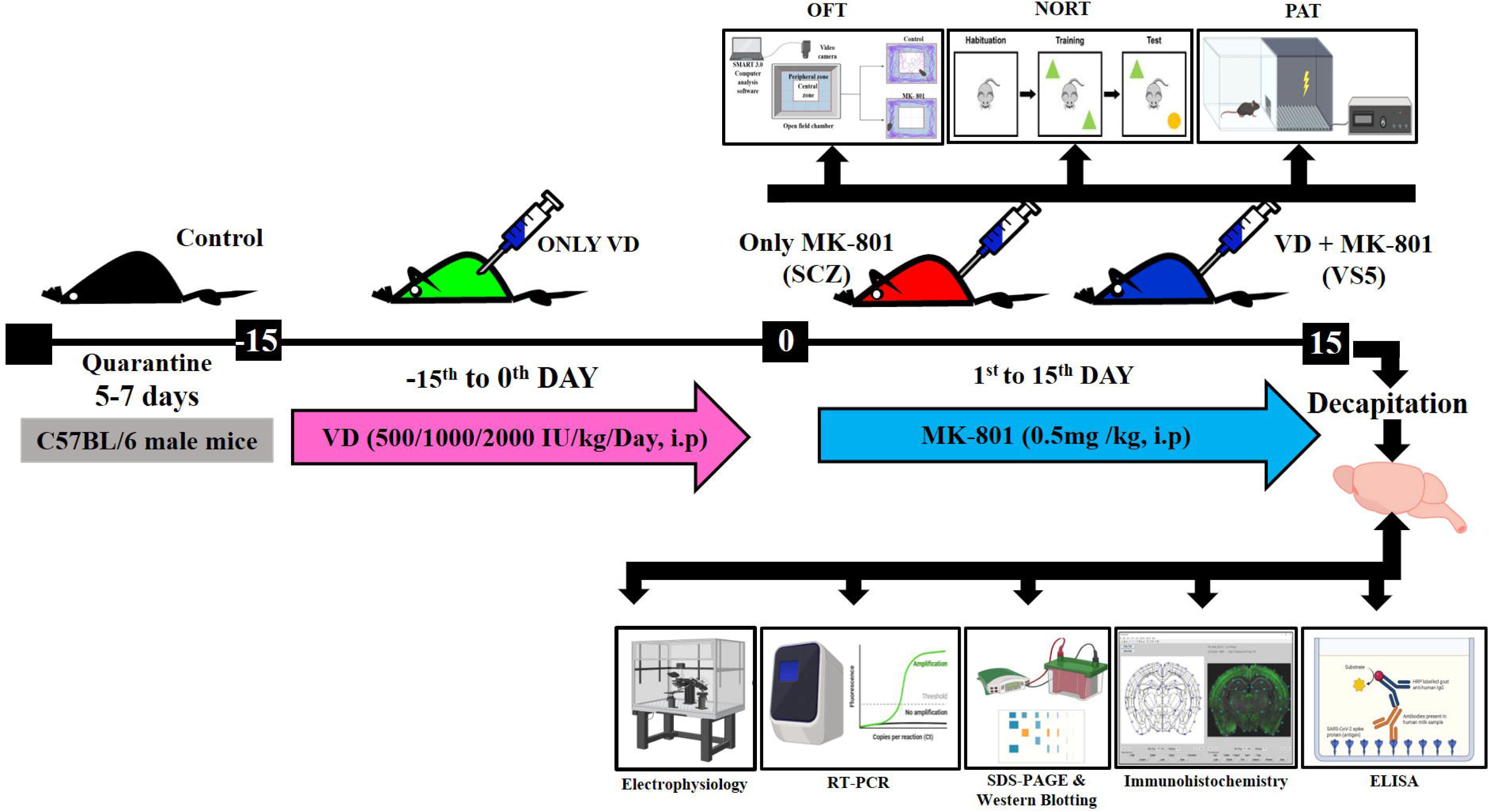
Design and timeline of the behavioral study. C57BL/6 male mice (8-10-week-old) were kept in quarantine for 5-7 days. VD5 and VS5 mice were pre-supplemented with 500IU/kg of VD for 15 days (Day −15 to Day 0). MK-801 was then injected at a dose of 0.5mg/kg to SCZ and VS5 mice after pre-supplementing with VD for 15 days (Day 0 to Day 15). Behavioral analysis were performed on Day 0, Day 7 and Day 15. On the 15^th^ day, mice were sacrificed and the PFC tissues were extracted for subsequent analysis.

### Drugs and reagents

a. **Cholecalciferol (Vitamin D3; VD)** was obtained (Sigma-Aldrich, India, Cat no.# C9756-1G) and dissolved in ethanol. Dilutions were prepared in sterile saline (final ethanol concentrations not exceeding cytotoxic levels of ≤0.1%), freshly made on each day of injection. **VD, VS5, VS1, VS2** groups of mice received VD (i.p) injections daily from −15^th^ to 0^th^ day as described previously [45].
b. **MK-801 hydrogen maleate (Dizocilpine)** (Sigma Aldrich, USA, Cat no.# M107-50MG) was dissolved in 0.9% saline to get a final concentration of 0.5 mg/kg. **SCZ**, **VS5**, **VS1**, and **VS2** groups of mice received MK-801 (i.p) injections daily for 15 consecutive days [72].

### Behavioral assessments

A total of almost 80 mice were used for the behavioral experiments. Before each experiment, mice were placed in the experimental room for 30 minutes to acclimatize to the surroundings (Fig. 1). Open field test was performed on the 1^st^, 7^th^ and the 15^th^ day whereas the novel object recognition test was performed on the 14^th^ day. Additionally, the passive avoidance test was performed on the 15^th^ day of post MK-801 administration (Fig. 1), where the training (shock) phase of passive avoidance test was performed only 4 hours before the test phase to minimize mice suffering.

### Assessment of locomotor activity using Open field test (OFT)

The open-field test was used to evaluate the locomotor activity of all groups of mice (i.e.**, Control, SCZ, VD5, VS5, VD1, VS1, VD2, and VS2**). The behavioral assay was conducted the 1^st^, 7^th^ and the 15^th^ day, 15 minutes after the MK-801 injection (via i.p at a dose of 0.5 mg/kg). MK-801 is known to produce hyperlocomotion, ataxia, staggering gait, and falling at a dose of 0.5 mg/kg [76]. MK-801 was administered once a day for 15 consecutive days and induces SCZ like symptoms. The OFT apparatus consisted of a white acrylic plexiglass square open area box (length 85 cm x width 75 cm x height 30 cm). An overhead camera attached to a computer (installed with SMART v.3.0 analysis software) tracked the trajectories and distance traveled by the mice. After allowing an initial 1 minute for adaptation to the box area, the total distance traveled during an additional 5 minutes was recorded as the locomotion activity [77]. After this, the mouse was placed back into its home cage, and the apparatus was cleaned with 70% ethanol to remove feces and allowed to dry completely before placing the next one. Each mouse was gently placed into the central zone of the box during the recording of its locomotion activity (Fig. 2). The total distance traveled in the center/periphery was analyzed using an automatic video tracking system (Smart version 2.5; Panlab, S.L.U., Barcelona, Spain) [77].

**Fig. 2.**
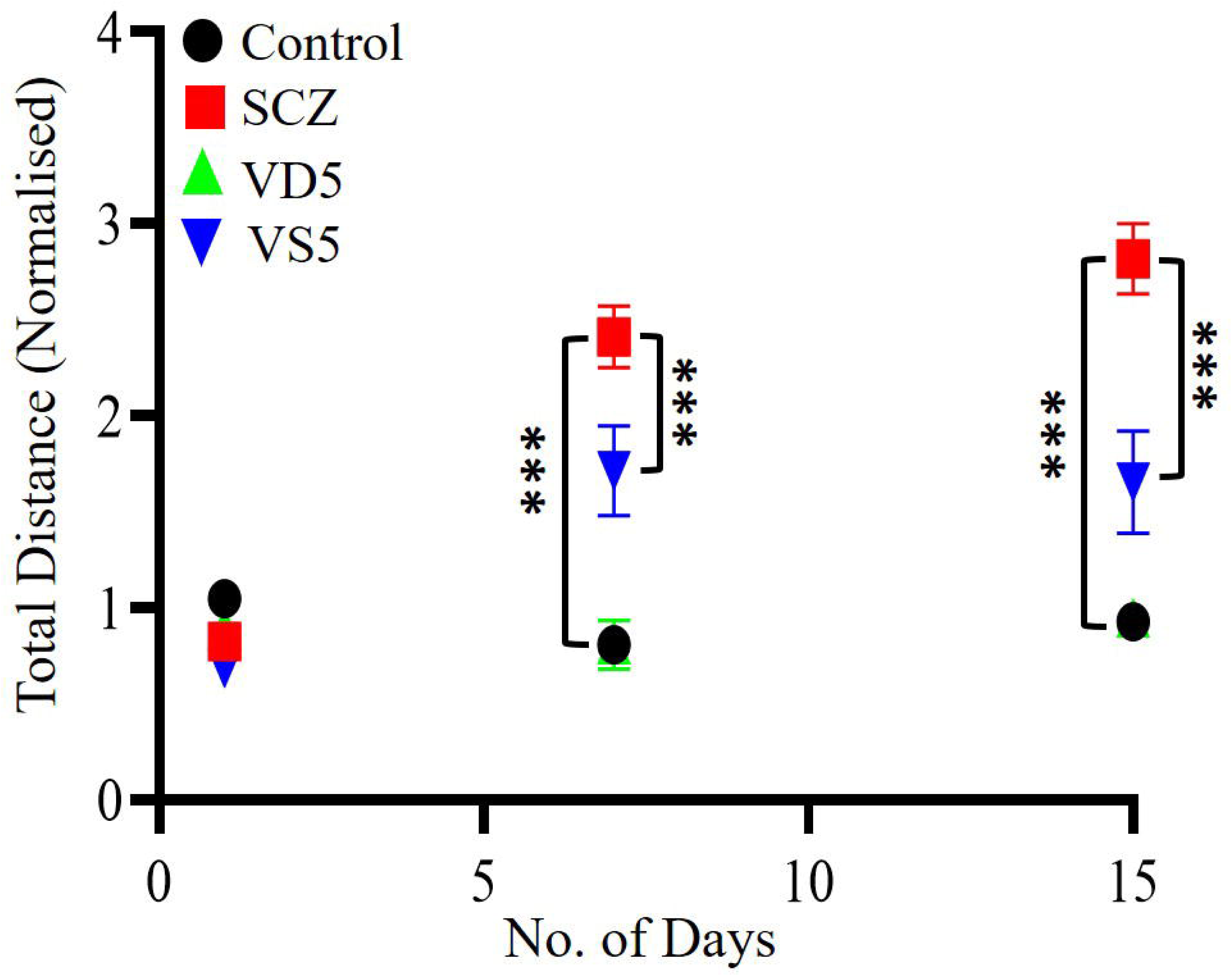
Pre-supplementation of VD at 500IU/kg alleviates hyperlocomotion in MK-801 induced SCZ mice. A highly significant difference was observed in the total distance travelled among the four groups, namely, **Control, SCZ, VD5 and VS5** (Two-way repeated measures ANOVA, group effect: F=52.42, p<0.001, Power = 1.00; day effect: F=88.28, p<0.001, Power=0.99). VS5 mice showed a significant reduction in the hyperlocomotion. A strong group and day interaction (f _(6)_ =32.09, p<0.001) indicated a beneficial effect of VD towards the alleviation of enhanced locomotory effect in SCZ mice.

### Evaluating learning and memory using Novel object recognition test (NORT)

The NOR test was performed as previously described with minor modifications [78]. It was conducted only on the 14^th^ day for the given groups, namely, **Control**, **SCZ**, **VD5**, and **VS5** mice. On the 13^th^ day, the habituation phase was undertaken, where the mouse was allowed to explore the test area (white, acrylic plexiglass box) for 5 minutes to avoid freezing behavior. On the next day, mice underwent 5 minutes of training, and after 30 minutes the test phase was performed. During the training phase, each mouse was exposed to two identical objects (a1 and a2; of the same size, shape, and color, proportionate to the size of the mouse (Fig. 3). These identical objects were placed diagonally to two opposite corners of the box. The mouse was allowed to explore both objects for 5 minutes, and the amount of time spent around each object was assessed using stopwatches. After 30 minutes post-training phase, the test phase was conducted where the mice were challenged with one familiar object (a1 or a2) and a new object (b) of different colors, shapes, and sizes. The testing phase was performed for 5 minutes. The box was cleaned before and after each trial with 70% ethanol before placing the next mice. Videos of each trial were taken using SMART v.3.0 software and were later analyzed by using stopwatches. The relative discrimination index was calculated to assess whether cognitive and working memory deficits impairment was induced by MK-801[78].

**Fig. 3.**
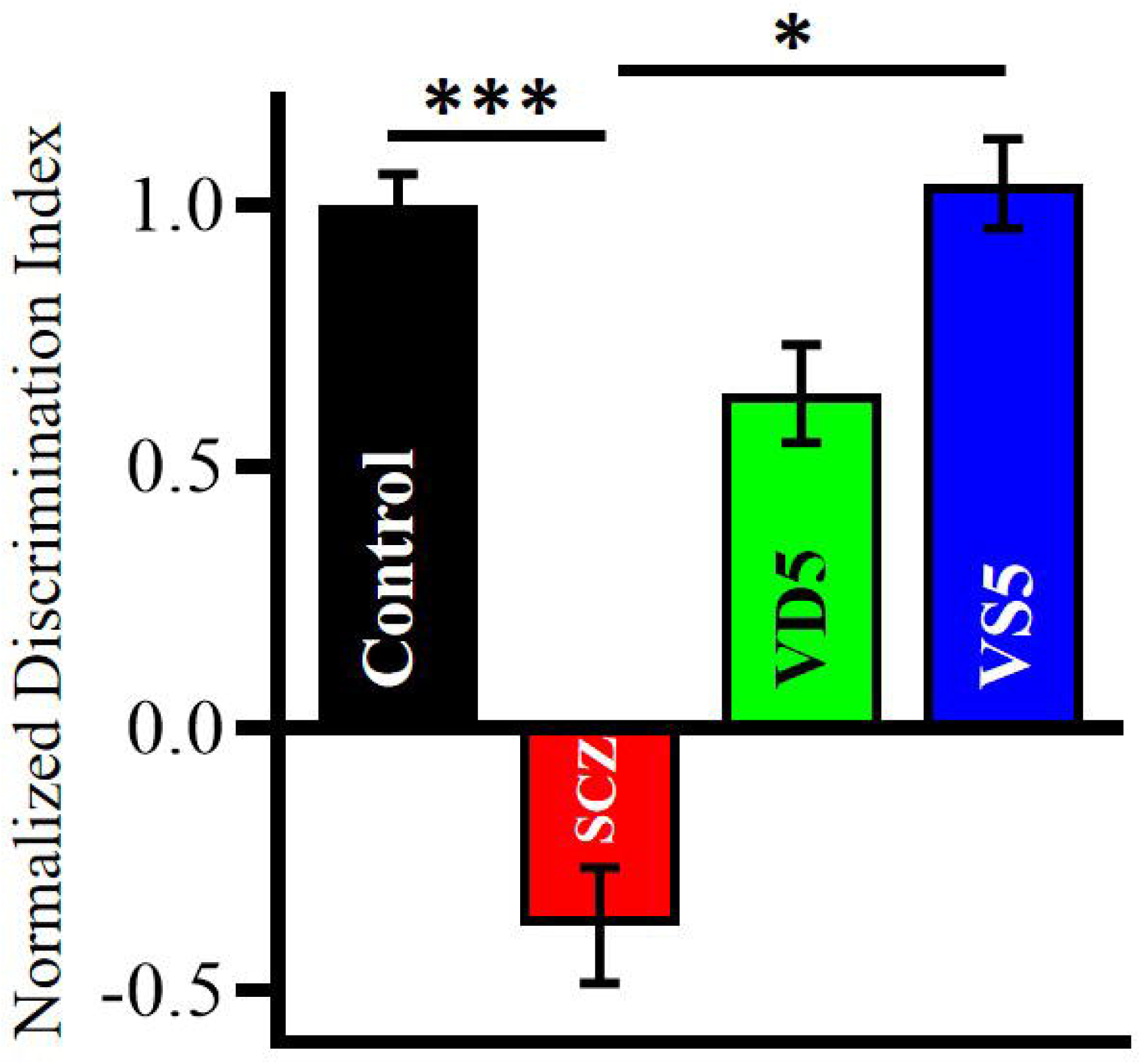
VD pre-supplementation rescues cognitive deficits in SCZ mice. MK-801 treatment significantly impaired recognition memory in SCZ mice compared to control (Control vs SCZ; n=12-14, p=0.0005, Student’s pairwise t-test), as evidenced by reduced preference for the novel object. This effect was substantially reversed by VD pre-supplementation on day 15 (SCZ vs VS5; n=12-14, p=0.015, Student’s pairwise t-test). Data is normalized, and represented as mean ± SEM.

### Passive avoidance test (PAT) for the assessment of short-term memory function

Passive Avoidance Test (PAT) was performed on the 15^th^ day to assess cognitive deficits associated with SCZ. The PAT apparatus included a closed compartment consisting of two acrylic boxes (a large white, roofless box and a small, black closed box provided with a detachable lid) connected through a guillotine door that shuts on the command of an analyzing software downloaded (ShutAvoid Software) to a computer. The boxes consisted of a detachable metal grill floor. The entry into the dark box was punished with a foot shock of 0.2 mA for 2 seconds. Each mouse was first allowed to explore the PAT apparatus for 5 minutes on the exploration day (24 hours before training) [79]. On the following day, ‘pre-test’ or ‘training’ was conducted. Each mouse was placed in the white, large box and given a waiting period of 30 seconds after which the guillotine door was opened, allowing access to the dark box. A trial period of 5 minutes (300 seconds) was conducted in which the mouse was punished with a foot shock upon entry into the dark box. For analysis of short-term memory, the ‘test’ was performed 4 hours after the training. No foot shock was given during the test. The time the mouse took to re-enter the dark box in 5 minutes during the test was considered as “latency”. The ‘test’ was done on the 15^th^ day of MK-801 injection for all the said groups, namely, **Control**, **SCZ**, **VD5**, and **VS5** mice respectively to assess memory function (Fig. 4).

**Fig. 4.**
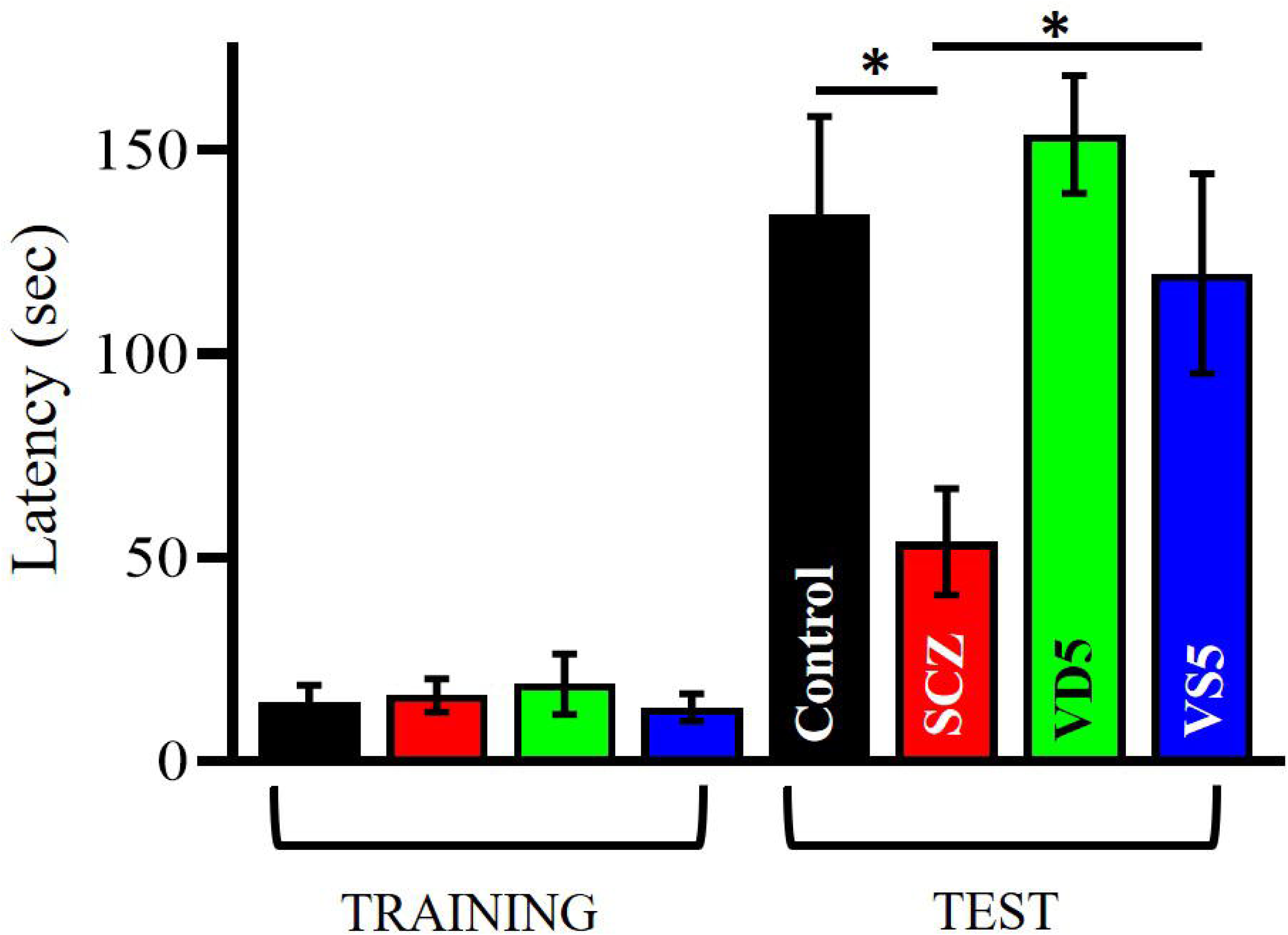
VD pre-supplementation rescue memory deficits in SCZ mice. On the 15^th^ day, VD intervention significantly ameliorated MK-801-induced memory deficits, as reflected by a reduction in latency in SCZ mice during the test phase (Control vs SCZ; p=0.028; SCZ vs VS5; n=13-17, p=0.02, Student’s pairwise t-test). Data is normalized, and represented as mean ± SEM.

### RNA isolation and cDNA preparation

Mice from the four groups (**Control, SCZ, VD5,** and **VS5**) post 15^th^ day of MK-801 administration, were anesthetized using ethyl ether (Sisco Research Laboratories, Cat no.# 64665) and immediately decapitated to extract the prefrontal cortex (PFC) brain tissue samples. The samples were then snap-frozen in liquid N_2_ and stored at −80°C until further use. To isolate RNA, the PFC brain tissue sample was homogenized in 1ml of RNAiso PLUS (Trizol, Takara Bio, Cat. # 9109) using a sonicator (ON-15 sec, OFF-10 sec; Total time-1 minute; 40% amplitude) as described previously [45]. Briefly, tissue samples were centrifuged at 14000 rpm at 4 °C (Eppendorf Refrigerated centrifuge, 542R) for 30 minutes after the addition of 200μl of chilled chloroform (Sisco Research Laboratories, Cat no.# 96764). RNA was precipitated from the aqueous layer with isopropanol (Hi-Media Laboratories, India, Cat no.# 62986), followed by an overnight incubation at −20 °C. Next day, the samples were centrifuged for 30 minutes at 4 °C at 14,000 rpm. They were washed twice with 70% chilled ethanol (Sisco Research Laboratories, Cas no.# 22072090) and centrifuged for 10 minutes at 4 °C at 14,000 rpm. Following two ethanol washes, the obtained pellet was resuspended in nuclease free water. RNA purity and concentration were determined using a nanodrop (Thermo Fisher Scientific, USA). RNA was then treated with DNase (Thermo Scientific, USA, Cat no.# EN052) to remove DNA contamination and the RNA concentration was rechecked using the nanodrop. Equal amounts of RNA from each group was used to reverse transcribe complementary DNA (cDNA) with the help of Verso cDNA synthesis kit (Thermo Scientific, USA, Cat no.# AB1453A) as per the manufacturer’s protocol. 1000ng of RNA was then taken from each group for cDNA synthesis with the following reaction conditions - 42°C for 1 hour and 95°C for 2 minutes. The resultant cDNA was used for real time polymerase chain reaction (RT-PCR) [45, 47].

### Quantitative expression analysis for genes by Real time PCR (RT-PCR)

The gene expression across all the four groups of mice (i.e. **Control, SCZ, VD5**, and **VS5**) from PFC brain tissue samples was analyzed on a CFX Opus 96 Real time PCR machine with the GoTaq qPCR SYBR mix (Promega Corporation, Cat no.# A6001). The reaction mixture was prepared as per the manufacturer’s recommended protocol using approximately 12ng of cDNA. The relative gene expression was quantified using the ΔCt method with specific primers (Table. 1) and then normalized to the 18s housekeeping gene. Fold changes in the target genes (NR1, NR2A, NR2B, NGF, BDNF, and α7nAChRs) was calculated using the 2^-ΔΔCt method with the Ct (Threshold cycle) values being obtained from the Bio-Rad CFX Manager 3.1 software and the relative fold change was calculated as follows - fold change = 2^−ΔΔCt, where ΔCt (cycle difference) = Ct (target gene) – Ct (Control gene) and ΔΔCt = ΔCt (treated condition) – ΔCt (Control condition) (Fig. 5 and 11)[45, 47].

**Fig. 5.**
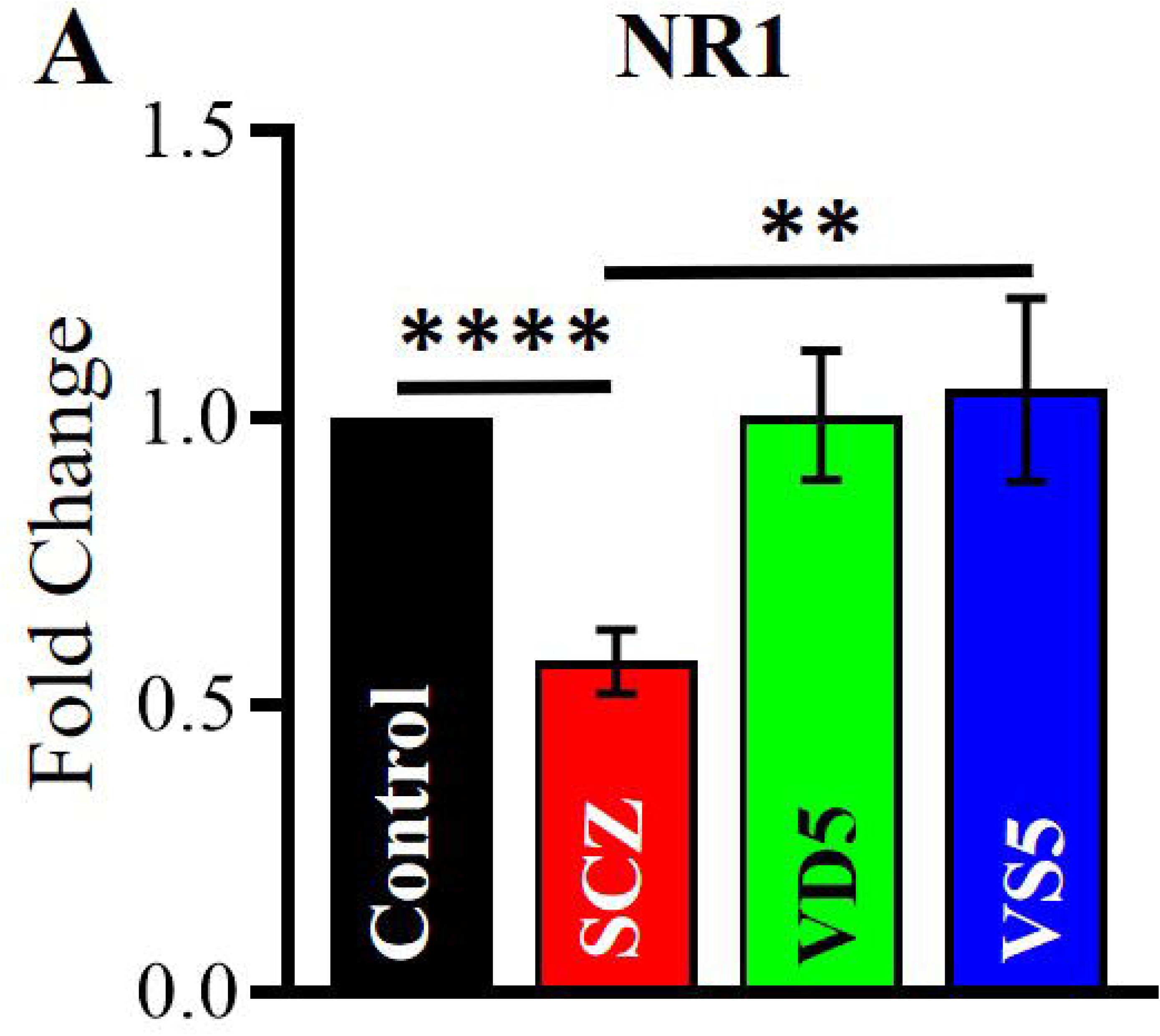
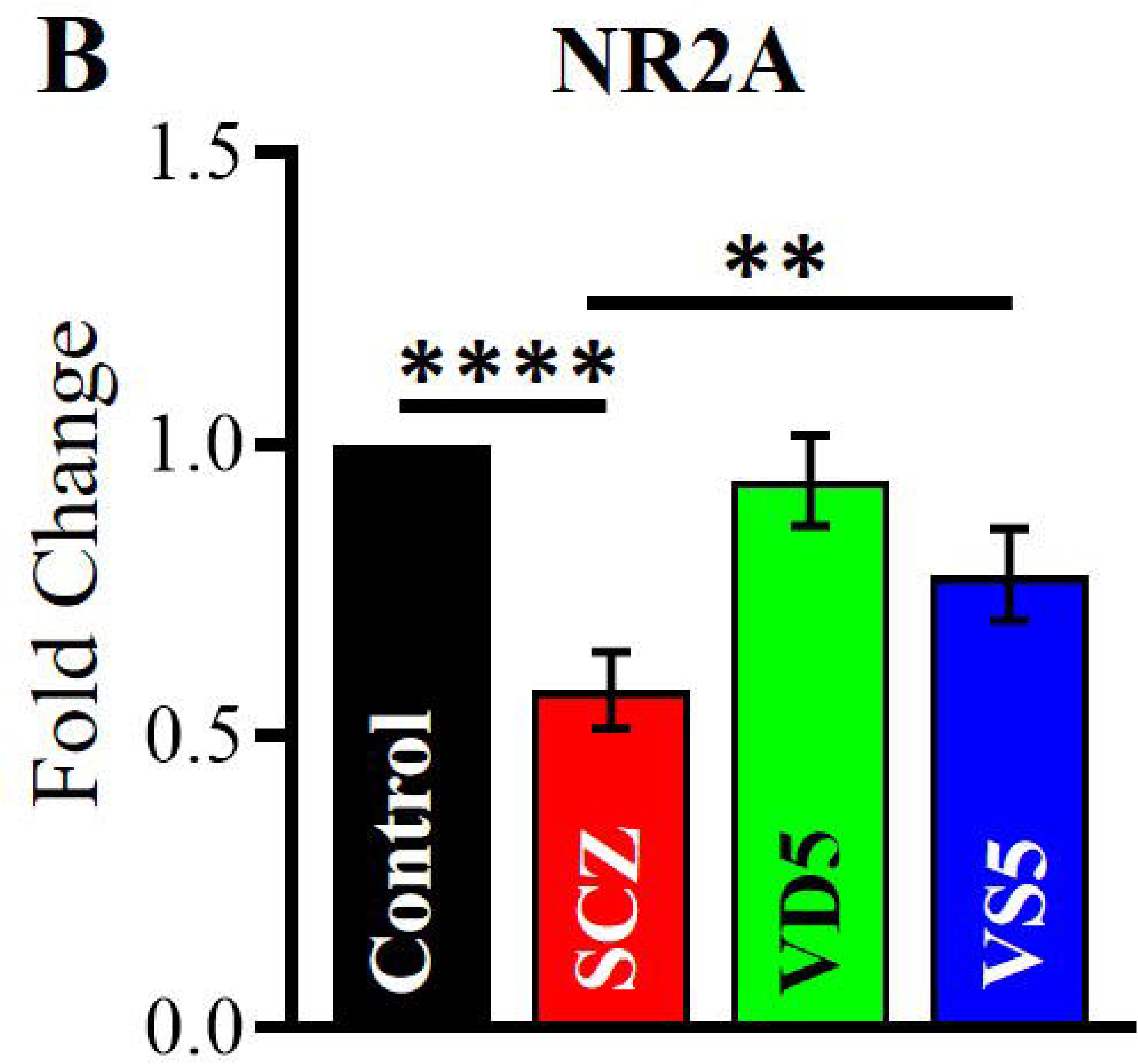
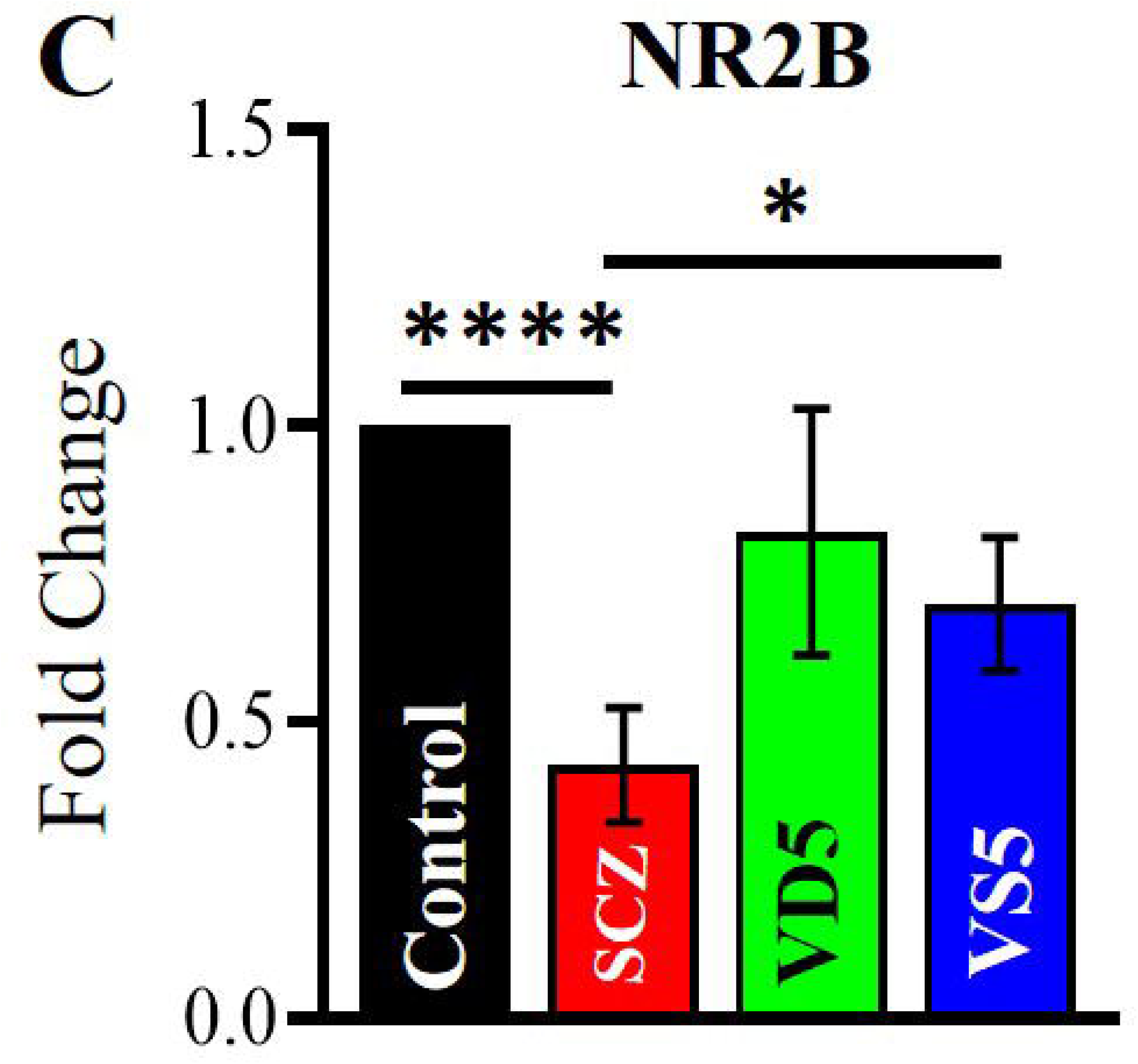
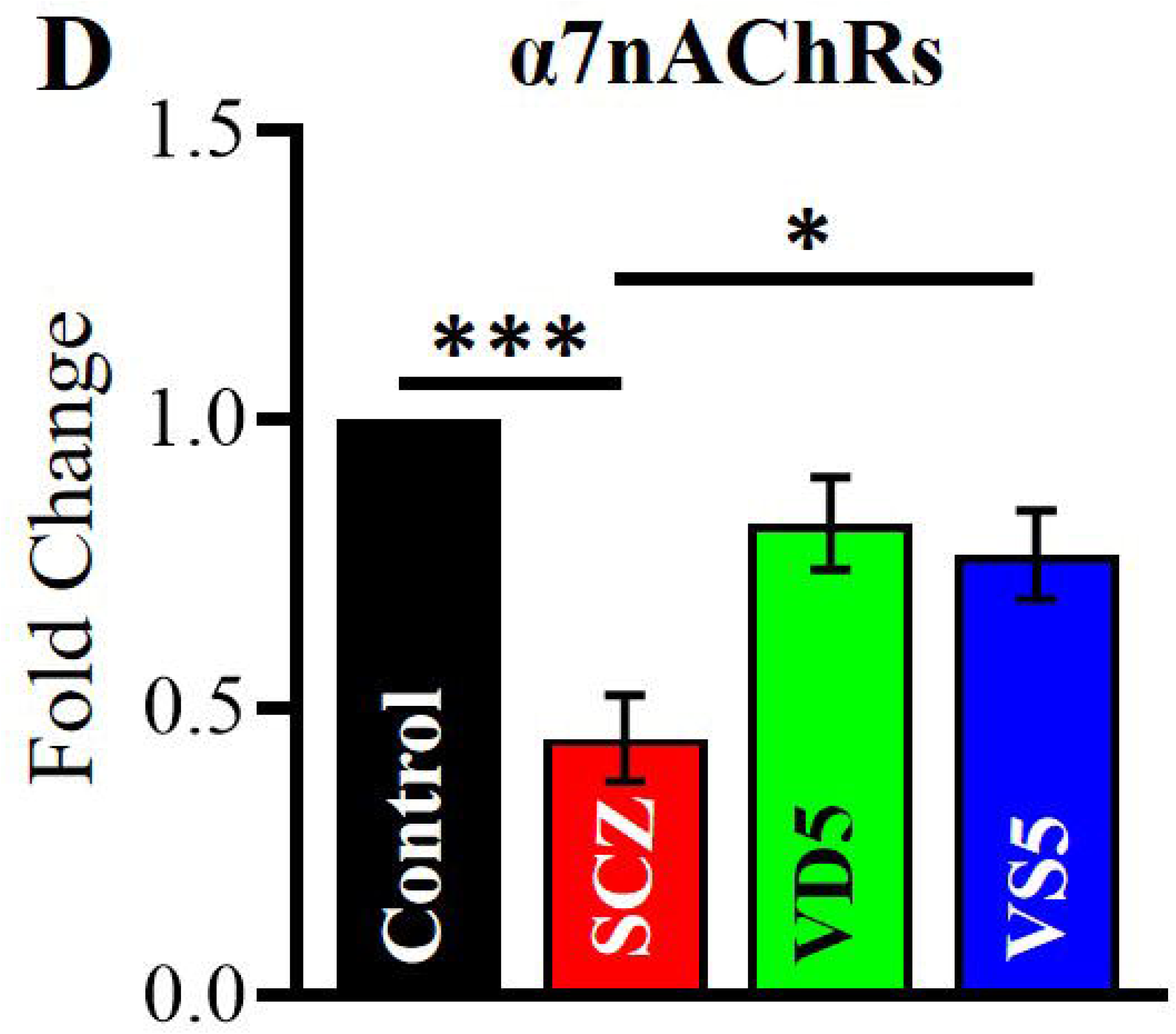
VD pre-supplementation rescues excitatory ionotropic receptor expression in the PFC of SCZ mice. **(A)** mRNA expression of NR1 was significantly reduced in SCZ mice (Control vs SCZ; n=6, p<0.0001, Student’s pairwise t-test) and was restored on VD presupplementation (SCZ vs VS5; n=6, p=0.005, Student’s pairwise t-test). **(B)** VS5 mice administered with VD showed a significant increase in the gene expression of NR2A (SCZ vs VS5; n=6, p<0.0001, Student’s pairwise t-test) which was downregulated in SCZ mice injected with MK-801 (Control vs SCZ; n=6, p=0.003, Student’s pairwise t-test). **(C)** A reduction in the mRNA expression of highly calcium permeable NMDA receptor, NR2B subunit was observed in the PFC (Control vs SCZ; n=6, p<0.0001, Student’s pairwise t-test) that got normalized in the VS5 mice (SCZ vs VS5; n=6, p=0.02, Student’s pairwise t-test). **(D)** A similar decrease in the mRNA expression of α7nAChRs was observed in the PFC (Control vs SCZ; n=4, p=0.0002, Student’s pairwise t-test) and got rescued when pre-supplemented with VD for 15 days (SCZ vs VS5; n=4, p=0.02, Student’s pairwise t-test). Data is normalized, and represented as mean ± SEM.

### Immunohistochemical analysis of α7nAChRs in the PFC

After the 15^th^ day of MK-801 administration, mice from four groups (**Control, SCZ, VD5, and VS5**) were deeply anesthetized using ethyl ether (Sisco Research Laboratories, Cat no.# 64665) and perfused with phosphate buffered saline (PBS) followed by 4% paraformaldehyde (PFA, Sigma Aldrich, Cat no.# 158127). The brains were post-fixed and stored in 4% paraformaldehyde (PFA) for 24-48 hours at 4°C, washed in PBS and cryoprotected in increasing concentrations of sucrose/glycerol (20-30%) until sectioning. 25µm brain sections were cut using a cryostat (Leica GM 1510S) and further mounted onto Super Frost glass slides as described previously [80]. The slides were air dried for 1-2 days and stored at −20°C for further use. For immunohistochemical studies, the slides were initially washed with 100% xylene (Finar, Cat no.# 1330-20-7) for two times and then rehydrated through a series of graded ethanol washes (100-50%), each wash of five minutes. This was followed by two PBS washes and an optional MilliQ wash. The slides were then subjected to antigen retrieval using boiling citrate buffer (10mM, pH 6.0, 100°C) (HiMedia, Cat no.# ML089) for 10 minutes. To reduce background fluorescence, the sections were treated with sodium borohydride (0.1% in PBS) (Biomatik, Cat no.# A3913) for five minutes. Further, permeabilization was performed using 0.3% Triton X (HiMedia, Cat no.# MB031) diluted in PBS for 40 minutes and blocked with 4% horse serum containing (Lonza, Cat no.# 14-403E)1% BSA for 2 hours at RT in a humid chamber. The slides were then incubated overnight at 4°C with α7nAChRs primary antibody (1:200) (Invitrogen, Cat no.# MA5-31691). The next day, the sections were washed with PBST and incubated with Anti-mouse secondary antibody (Anti-mouse IgG (H+L), F(ab’)2 Fragment; Alexa Fluor® 488 Conjugate, Cell signaling technology, Cat no.# 4408) for 2 hours at RT in dark conditions. The final washes were followed by mounting the sections using DAPI (Invitrogen, Cat no.# S36964) containing mounting media. Imaging was performed using a confocal microscope (FLUOVIEW FV10i; Olympus) under identical acquisition settings for all the experimental groups. The images acquired were quantified using ImageJ (Fig. 6).

**Fig. 6.**
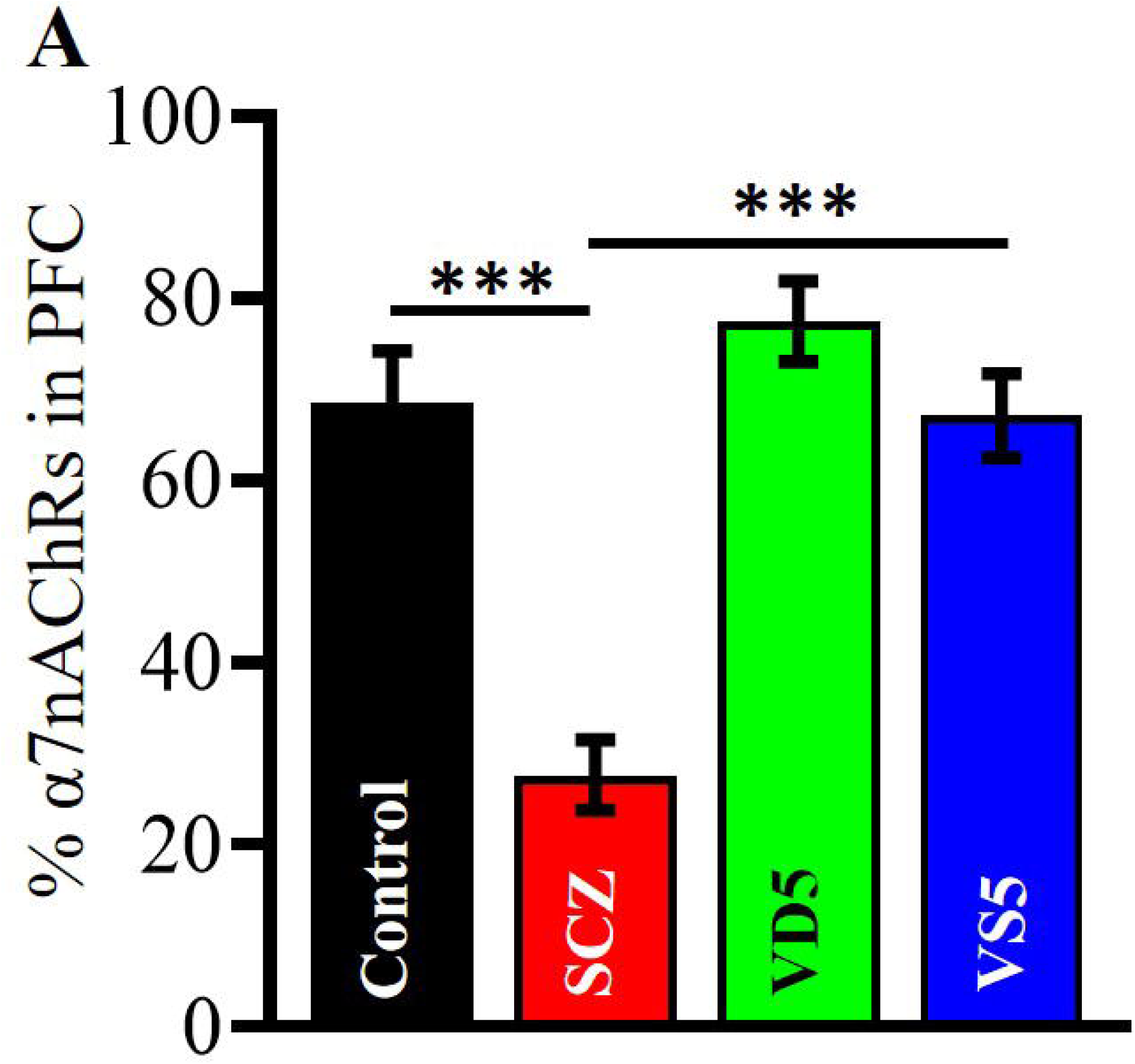
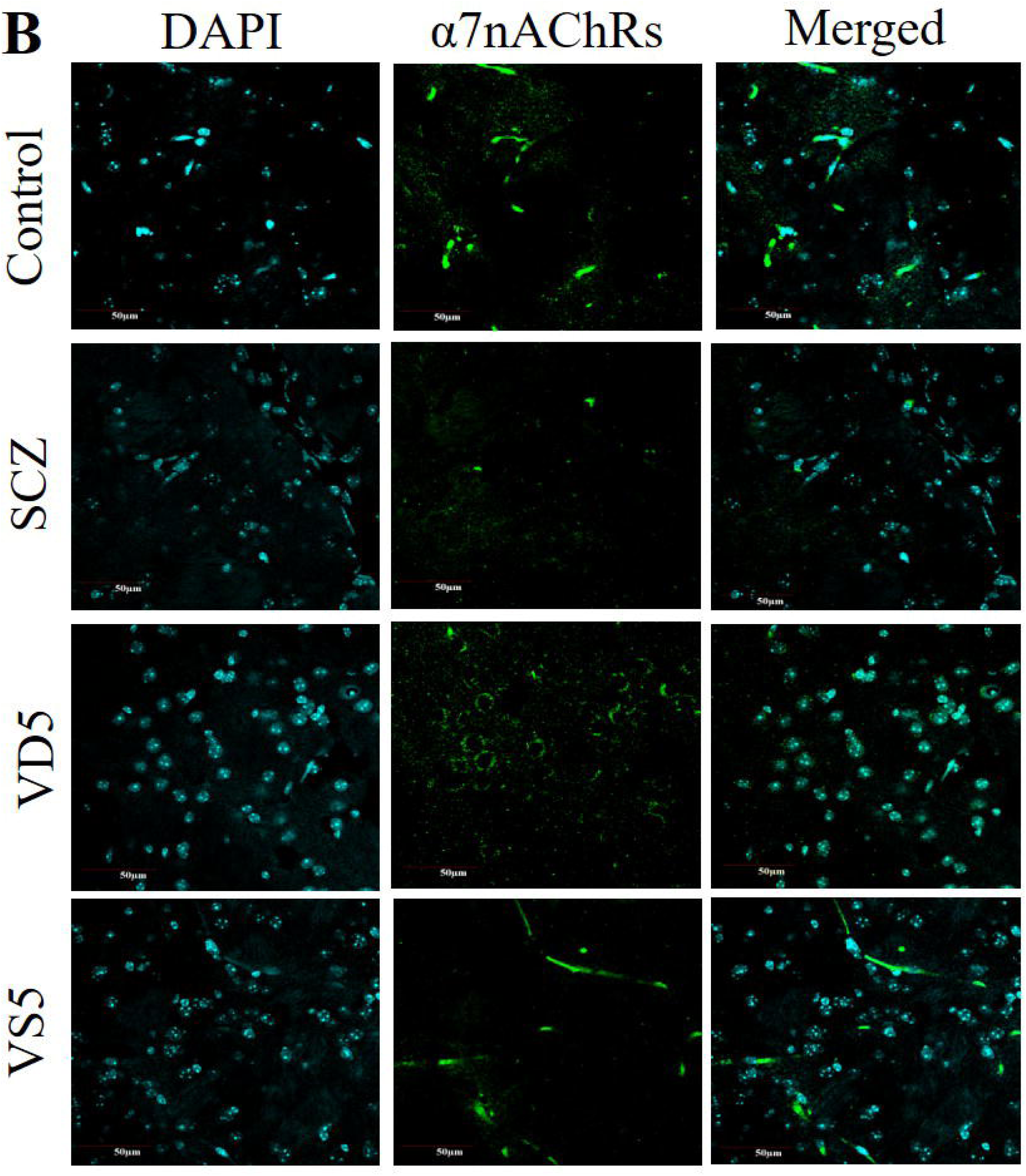
VD pre-supplementation restores α7nAChRs expression in the PFC of SCZ mice. (A) Immunohistochemical analysis of α7nAChRs in the PFC showed a marked reduction in the receptor immunoreactivity in SCZ samples when compared to age matched controls (Control vs SCZ; n=3 animals, F (3, 31) =23.4, p<0.0001, one-way ANOVA), characterized by diminished staining intensity and altered distribution across the various cortical layers. **(B)** Representative images demonstrating a significant restoration in the α7nAChRs protein expression in the PFC on VD pre-supplementation in SCZ mice (SCZ vs VS5; n=3 animals, F (3, 31) =23.4, p<0.0001, one-way ANOVA; Scale bar=50 µm) highlighting its neuroprotective role.

### Acetylcholinesterase (AChE) activity assay

The Acetylcholinesterase (AChE) activity across the four experimental groups **(Control, SCZ, VD5, and VS5)** was measured using the Amplex® Red Acetylcholine/Acetylcholinesterase Kit (Invitrogen, Cat no.# A-12217) in accordance with the protocol directed by the manufacturer. This assay provides an ultrasensitive method for detecting acetylcholinesterase in a fluorescence microplate reader. It indirectly assesses the AChE activity with the help of a highly sensitive fluorogenic probe for horseradish peroxidase (HRP) called Amplex red. In this assay, AChE first converts acetylcholine substrate to choline. Choline further gets oxidized into H_2_O_2_ and betaine using the choline oxidase enzyme. H_2_O_2_ in the presence of horseradish peroxidase reacts with Amplex red reagent (1:1 ratio) to give rise to a fluorescent resorufin product which is measured using a fluorescent plate reader (Spiromax, USA). To perform this assay, 100 μl of 1:10 diluted brain tissue sample from respective group of mice was incubated with 100 μl of working solution (200 μM Amplex red reagent, 1 U/ml horseradish peroxidase (HRP), 0.1U/ml Choline oxidase (CO), 50 μM Acetylcholine, Volume made up with 1X reaction buffer). The mixture was incubated for 30 minutes and the fluorescence intensity was measured at 540 excitation - 590 emission wavelengths. AChE activity was determined by plotting a standard curve and expressed as mU/mg protein after subtracting the background fluorescence values for each sample [47] (Fig. 7).

**Fig. 7.**
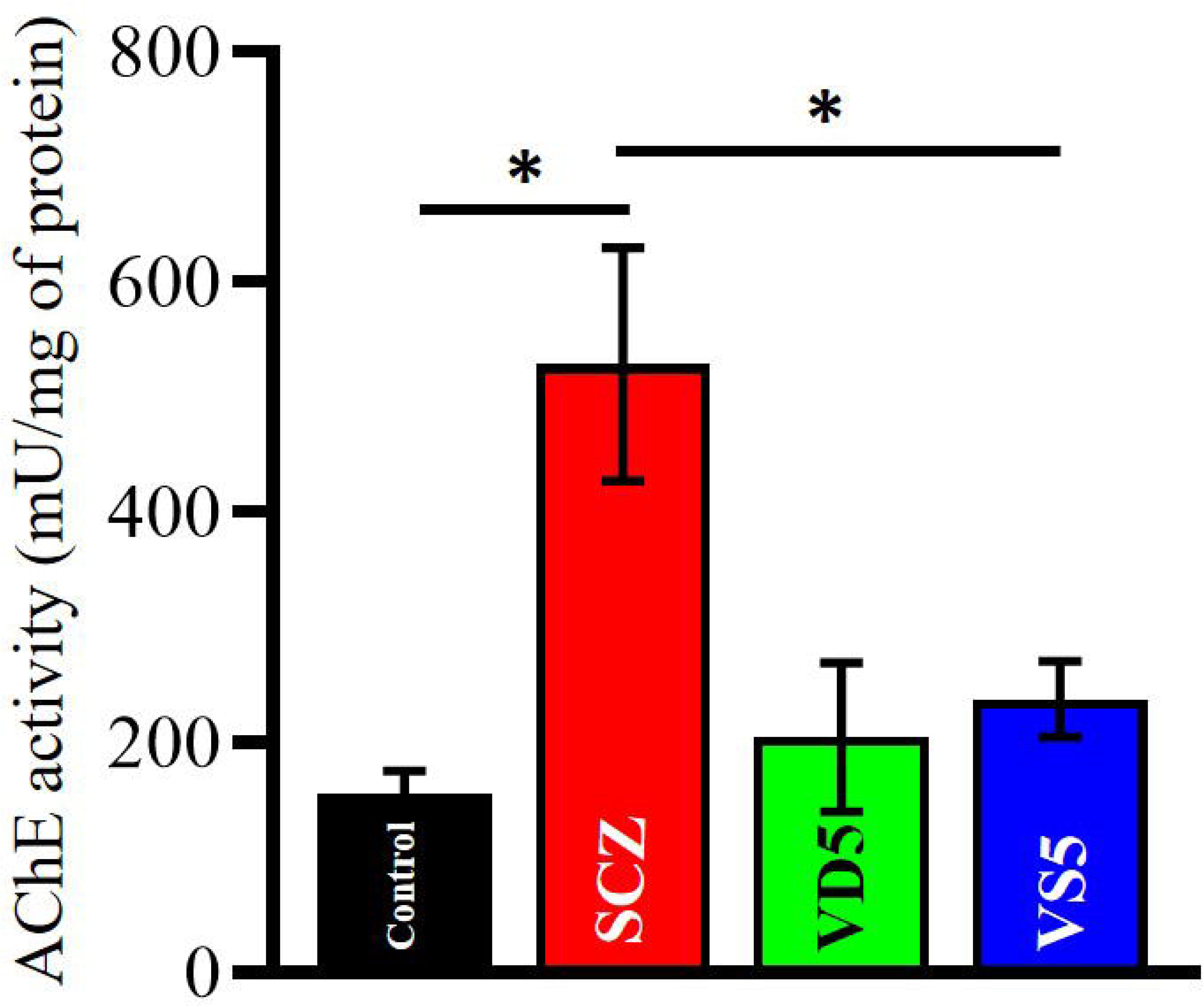
Pre-supplementation of VD alleviates acetylcholinesterase levels in the PFC of SCZ mice. SCZ mice showed an increment in AChE activity (Control vs SCZ; n=5, p=0.01, Student’s pairwise t-test) that got severely attenuated on VD intervention in VS5 mice (SCZ vs VS5; n=5, p=0.04, Student’s pairwise t-test). Data represented as mean ± SEM.

### Quantification of calcidiol levels

Blood was collected via the retro-orbital sinus from the four experimental groups (**Control, SCZ, VD5, and VS5**) on the 15^th^ day and transferred to pre-chilled EDTA-coated microtubes. It was centrifuged at 4000 rpm for 20 minutes at 4 °C to isolate plasma. The resultant supernatant was stored at −80 °C for further experiments. The plasma calcifediol (or 25-hydroxy Vitamin D3- [25(OH)D]) were quantified using the General 25 Hydroxy Vitamin D3 ELISA Kit (BT Lab, Cat no.# EA0005Ge) as per the manufacturer’s instructions. Plasma samples were added to the 96 well plate pre-coated with anti - 25(OH)D antibody, then incubated with HRP-conjugated VD and further developed with TMB substrate. The plate was read at an absorbance of 450 nm using a microplate reader (Spiromax, USA). The concentrations were then calculated by plotting a standard curve and were expressed as ug/ml of calcifediol after subtracting background absorbance values for each of the samples. Additionally, serum calcifediol/calcidiol (or 25-hydroxy Vitamin D3- [25(OH)D]) concentrations were also measured from patient samples (Control vs SCZ) using the CPC iFlash 1200 Chemiluminescence Immunoassay (CLIA) Analyzer (CPC Diagnostics Pvt.Ltd., Chennai, India). Venous blood (3-5ml) was drawn from each participant using standard aseptic precautions. Blood was allowed to clot at room temperature for half an hour, and centrifuged at 3000 rpm for 10 minutes for the collection of the serum. The supernatants obtained were stored at 2°C - 8°C, and processed within 48 hours of collection. Commercially available chemiluminescent immunoassay kit (CPC Diagnostics, Cat no.# C86023) was used to quantify the 25(OH)D (calcifediol) levels in guidance to the manufacturer’s protocol (Fig. 8).

**Fig. 8.**
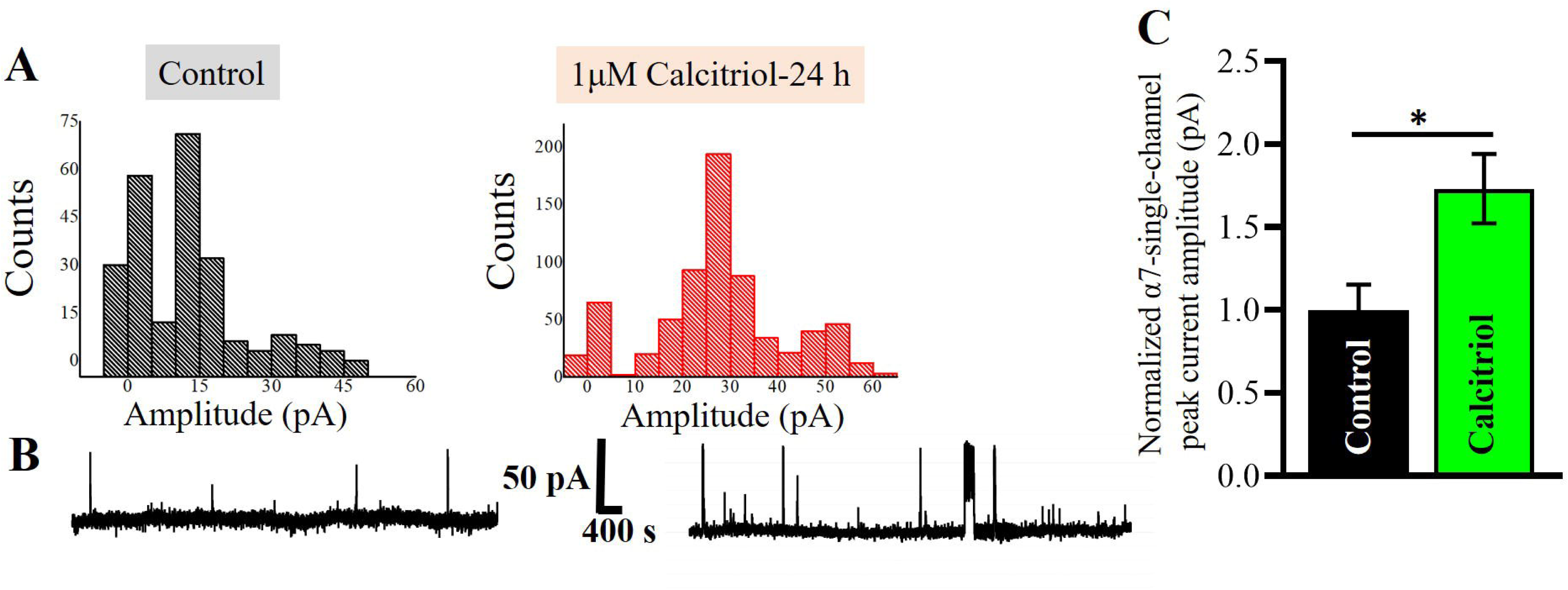
Hypovitaminosis in SCZ mice and Indian SCZ patients. **(A)** Circulating levels of calcidiol (25-hydroxyVitamin D3; ug/L) in mouse plasma were found to be lower in MK-801 induced SCZ samples as compared to controls (n=5). Data is represented as mean ± SEM. **(B)** Circulating levels of calcidiol (25-hydroxyVitamin D3; ng/L) in the serum were significantly reduced in SCZ patients (n=113) when compared to healthy controls (n=73) (Control vs SCZ; U=3299.5, p=0.021, Mann-Whitney U test).

### Cell-culture, transfection, and fluorescence imaging

Fluorescently tagged α7nAChRs with Venus fluorophore (α7-Venus) and RIC-3-CFP subunits were kindly provided by Professor Raad Nashmi’s Lab (University of Victoria, British Columbia, Canada) [81–83]. The plasmids were transformed into DH5–alpha competent cells. Transformation of the product into the bacterial cells was then confirmed by spreading it on LB agar plates containing ampicillin (100 µg/ml), which yielded individual colonies containing the plasmid constructs. A few colonies from the transformed plates were picked, inoculated, and plasmids were extracted the next day using the Qiagen Miniprep method (Qiagen, Cat. No.# 27104). The product obtained was then run on 1% agarose gel and subjected to electrophoresis to confirm the presence of the plasmid.

Human embryonic kidney HEK293T cells (ATCC) were maintained and cultured as done previously [84]. Briefly, these cells were grown using Dulbecco’s Modified Eagle’s medium (DMEM, Thermo Fisher Scientific, Cat no.# 11995065), supplemented with 10% Fetal Bovine Serum (FBS; Thermo Fisher Scientific, Cat no.# 16000044), 1% L-Glutamine (Thermo Fisher Scientific, Cat no.# 35050061), penicillin (100 μg/ml), and streptomycin (100 μg/ml) (Thermo Fisher Scientific, Cat no.# 15140122). The cells were transiently transfected with α7-Venus and RIC-3-CFP using Lipofectamine 1000 (Thermo Fisher Scientific, Cat no.# 18324010) transfection reagent. Equimolar concentrations of α7-Venus and RIC-3 CFP were mixed with Lipofectamine 1000 transfection reagent. 24 hours post-transfection, the cells were treated with 1μM calcitriol (Med chem Express, Cat no.# HY-10002) to record single-channel currents mediated by α7nAChRs [84].

### Single channel recording via cell-attached- patch-clamp electrophysiology

Single-channel recordings in the cell-attached patch-clamp configuration was performed at room temperature on HEK293T cells transfected with mouse α7-Venus tagged nAChRs and RIC-3 cDNA [84]. The bath and pipette solutions contained the following (in mM):150 NaCl,4 KCl, 2 CaCl2, 2 MgCl2, 10 HEPES, and 10 D-glucose, pH 7.4, with 100 µM nicotine dissolved in the pipette solution. Micropipette recording electrodes were 1.5mm OD and 1.0 mm ID borosilicate glass (DSS Imagetech, Cat no.# G-1.5) that were pulled on a narishige puller (PC 100; Serial no: 22108). The pulled pipettes were coated with Sylgard (Sigma Aldrich, Cat no.# 761036-5EA) and fire polished with a microforge (DSS Imagetech, Cat no.# MF2). Recording micropipettes had resistances between 8 and 12 MΩ and contained 1mM nicotine (Sigma, Cat no.# N3876). A pipette holding potential (Vp) of 40 mV was used throughout the recordings. Single-channel currents were acquired using a Multiclamp 700B amplifier (Molecular Devices), low-pass filtered at 4 kHz, digitized at 50kHz with a Digidata1550 converter (Molecular Devices), and collected with pClamp 11.3 software. Clampfit 11.3 software (Molecular Devices) that was used to determine single-channel amplitudes. Single-channel recordings were notch-filtered at 60 Hz, followed by a 4 kHz Gaussian filter. Single-channel current amplitudes were analyzed by single-channel events in Clampfit 11.3 software (Fig. 9).

**Fig. 9.**
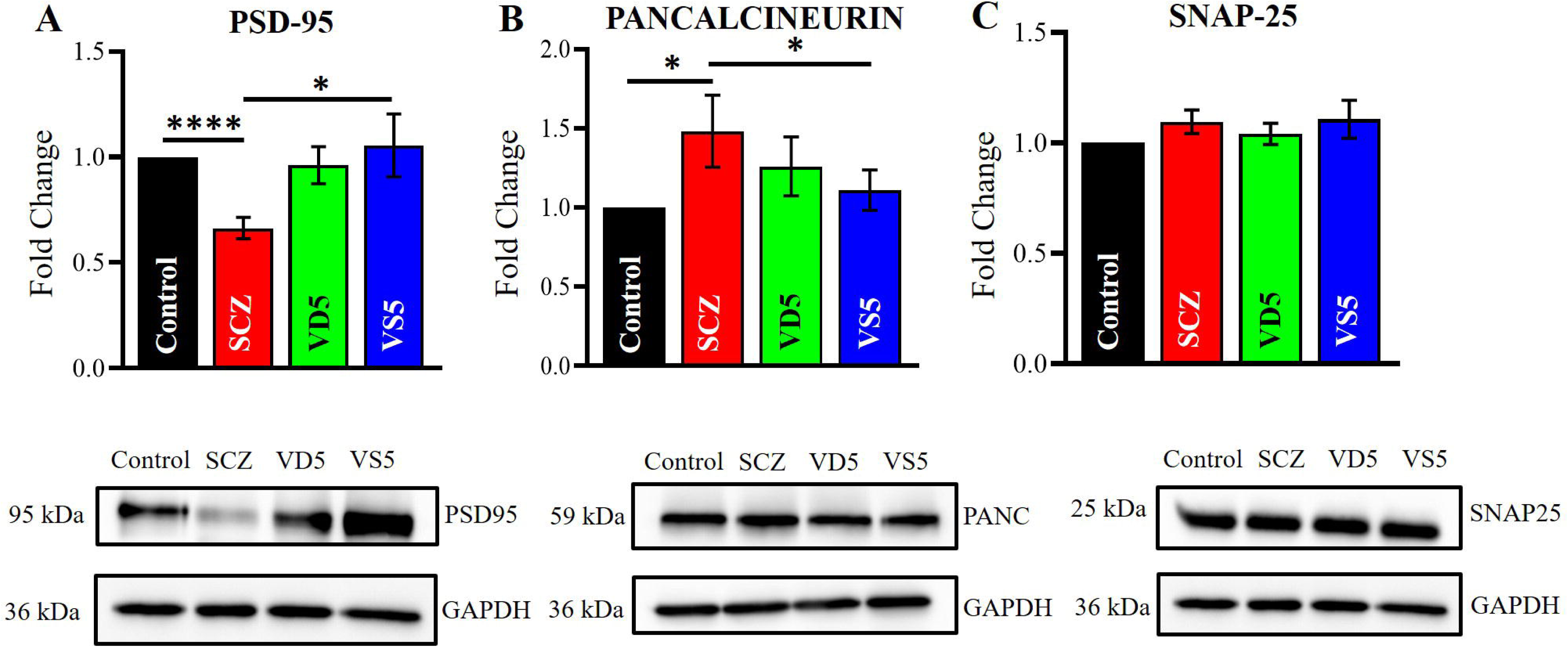
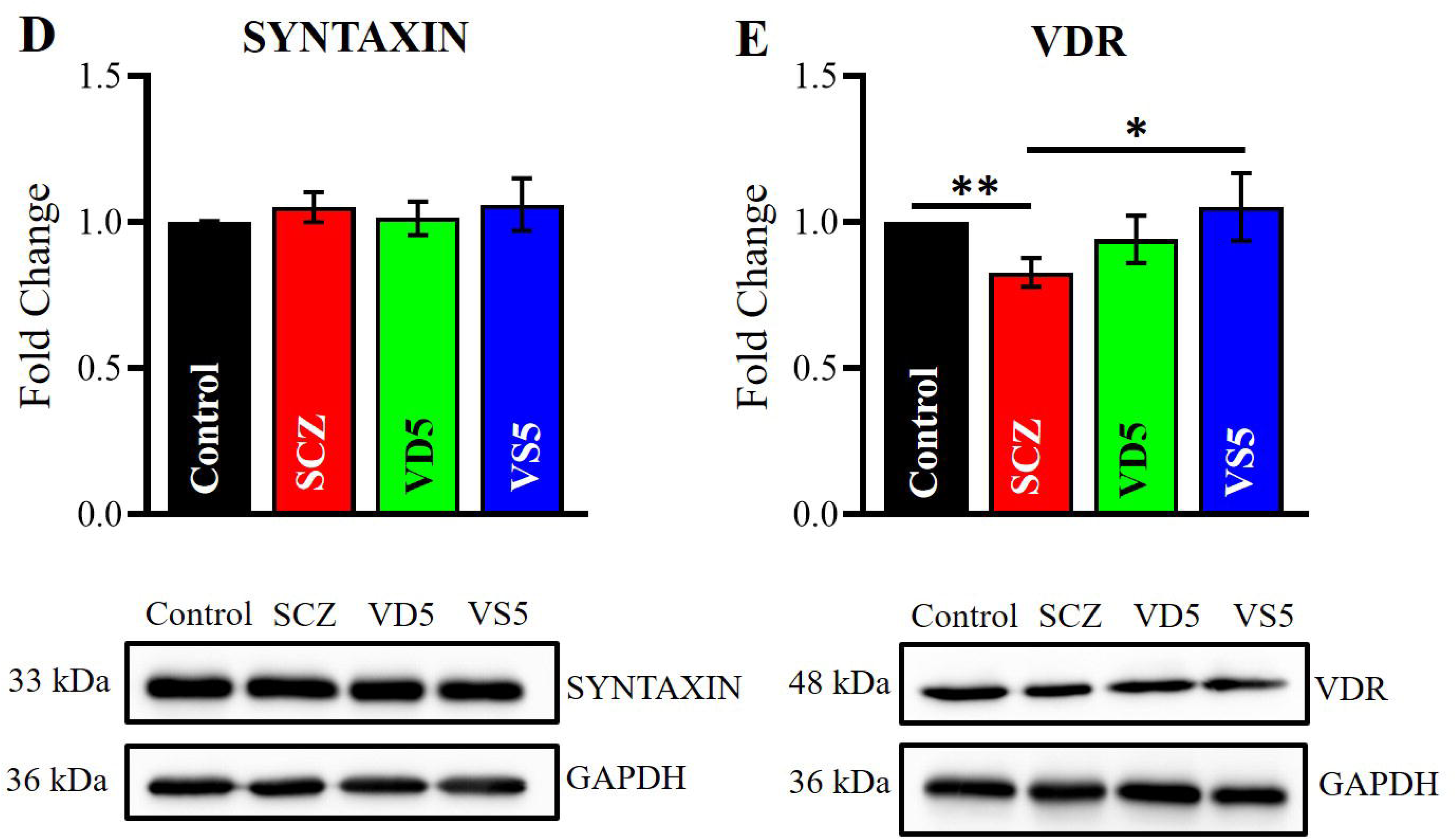
Calcitriol enhances α7nAChRs mediated single-channel currents in HEK293T cells. (A) Calcitriol (1μM) pretreatment for 24h significantly enhanced α7nAChRs single channel peak current amplitude as compared to control, with amplitude histograms showing rightward shift. **(B)** Representative traces demonstrating single-channel current events recorded in cell-attached configuration from HEK293T cells with 1mM of nicotine in the recording pipette. **(C)** Bar graph representating normalized mean peak single-channel current amplitude of control with and without calcitriol (Control vs Calcitriol; n=4, p<0.02, Student’s pairwise t-test). Data is normalized, and represented as mean ± SEM.

### Protein quantification and western blotting

To examine whether VD influences the protein expression of synaptic markers implicated in SCZ, various synaptic proteins were analyzed using SDS-PAGE and western blotting as described previously [45, 47]. Prefrontal cortex (PFC) brain tissue samples were extracted on the 15^th^ day from the four experimental groups (**Control, SCZ, VD5, and VS5**) of mice. The tissues were homogenized in RIPA lysis buffer (150mM sodium chloride, 50mM Tris; pH 8.0, 0.5% sodium dodecyl sulfate, 1% Triton X-100 and protease inhibitor cocktail (PIC)). Total protein was isolated and the concentrations were measured using the Bradford protein assay kit (Bio-Rad, Cat no.# 5000006). Equal amounts of protein (50 μg) were loaded and separated on a 12% SDS-PAGE gel and then transferred onto a PVDF membrane using a trans blot wet transfer system (Bio-Rad, USA). The membranes were blocked using 5% BSA (Sisco Research Laboratories, Cat no.# 85171) and further incubated with respective primary (Table. 2) and secondary antibody (Anti – rabbit IgG, HRP-linked Antibody, 1:5000) (Cell signaling technology, Cat no.# 7074). GAPDH (1:1000, Cell signaling technology, Cat no.# D16H11) served as a loading control for normalization. The protein bands were visualized using the fusion pulse gel documentation system (Eppendorf, USA) and quantified using the ImageJ software (Fig. 10).

**Fig. 10.**
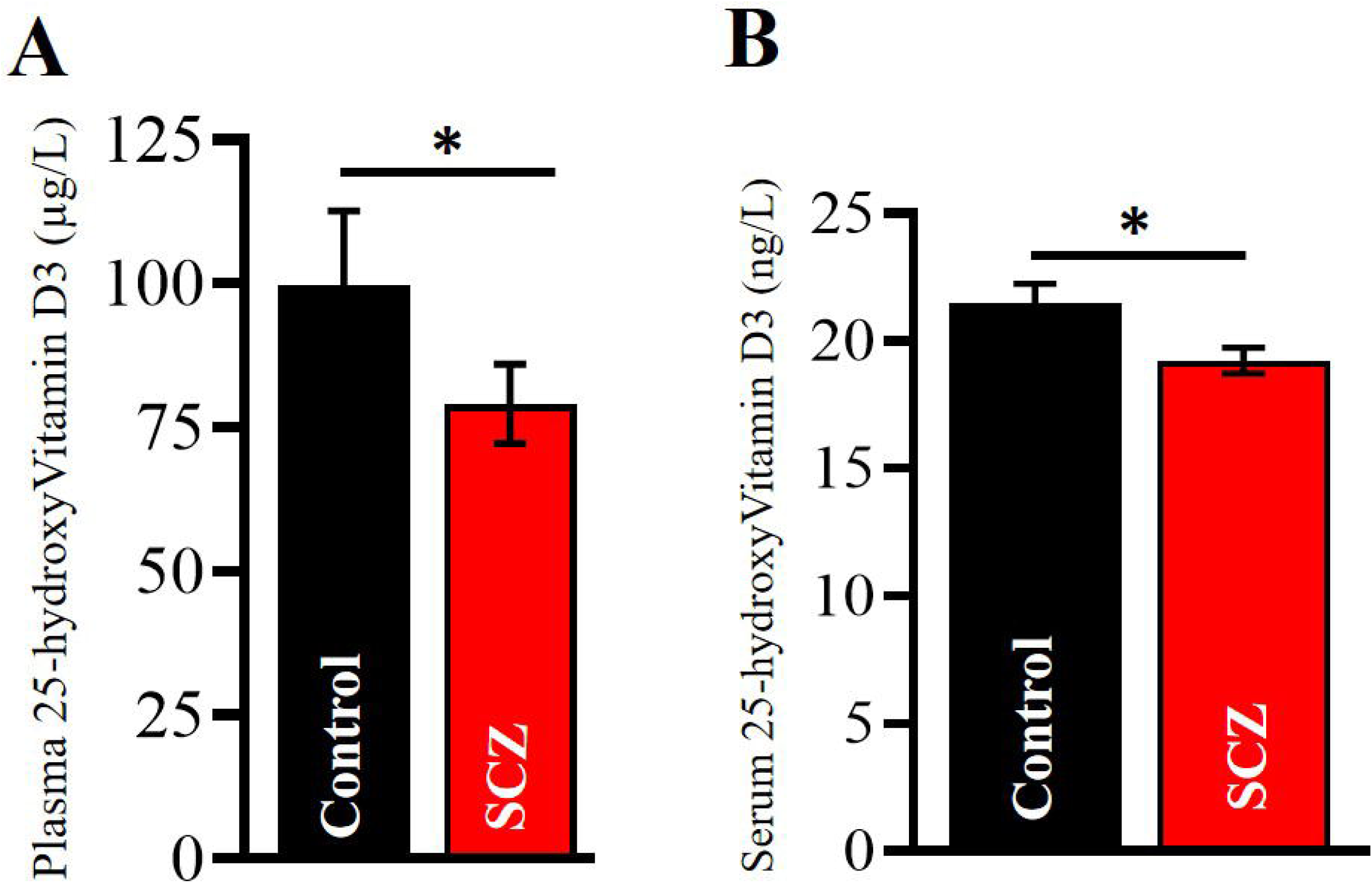
VD pre-supplementation restores synaptic protein expression in the PFC of SCZ mice. **(A)** The protein expression of PSD-95 was significantly decreased in SCZ mice (Control vs SCZ, n=4, p<0.0001, Student’s pairwise t-test). VD pre-supplementation significantly reversed its expression in VS5 mice (SCZ vs VS5; n=4, p=0.041, Student’s pairwise t-test). **(B)** Pan-calcineurin levels were highly increased in SCZ mice relative to controls (Control vs SCZ; n=6, p=0.04, Student’s pairwise t-test) and VD intervention in VS5 mice alleviated its protein expression (SCZ vs VS5; n=6, p=0.03, Student’s pairwise t-test). **(C, D)** Syntaxin and SNAP-25 protein expression remained unaltered. **(E)** VD pretreatment restored the VDR protein expression that was found be reduced in SCZ mice (Control vs SCZ; n=5, p=0.002; SCZ vs VS5; n=5, p=0.03, Student’s pairwise t-test). Data is normalized, and represented as mean ± SEM.

### *In-Silico* prediction of Vitamin D Response Elements (VDREs)

The putative Vitamin D Response Elements (VDREs) within the promoter regions of the selected genes were identified using the JASPAR transcription factor binding profile database. The promoter regions relative to the transcription start site (TSS) were extracted from the mouse genome using the coordinates obtained from the NCBI Gene database. For genes located on the positive strand, the promoter regions were defined as TSS −2000 bp to TSS +200 bp, whereas for genes located on the negative strand, coordinates were adjusted according to the strand orientation. The retrieved sequences were analyzed using the JASPAR CORE Vertebrates database by scanning for the binding motifs corresponding to the Vitamin D Receptor (VDR) or VDR: RXR heterodimer. The predicted binding sites were ranked based on their relative profile scores, and motifs with a relative profile score threshold of 75-80% located within the promoter regions were considered putative VDREs (Table. 3).

### TNF-α Quantification by ELISA

The levels of tumor necrosis factor-α were quantified using a commercially available ELISA kit (Highly Sensitive Enzyme Linked Immunosorbent assay (ELISA) Kit for Tumor Necrosis Factor Alpha (TNF-α); Cloud clone Corp, Cat no.# HEA133Mu). Pre-frontal cortex brain tissues from the four experimental groups (**Control, SCZ, VD5, and VS5**) were lysed in ice-cold RIPA buffer, centrifuged and were isolated for total protein. Total protein concentration was estimated using Bradford assay (Bio-Rad, Cat no.# 5000006). Standards and samples were added to the 96 well pre-coated plate in duplicates, incubated with biotin conjugated detection antibody and HRP-conjugated streptavidin and further were developed with TMB substrate. The plate was read at an absorbance of 450nm using a microplate reader (Spiromax, USA). The concentrations were then calculated by plotting a standard curve and were expressed as pg/ml of TNF-α after subtracting background absorbance values for each of the samples [47] (Fig. 12).

**Fig. 11.**
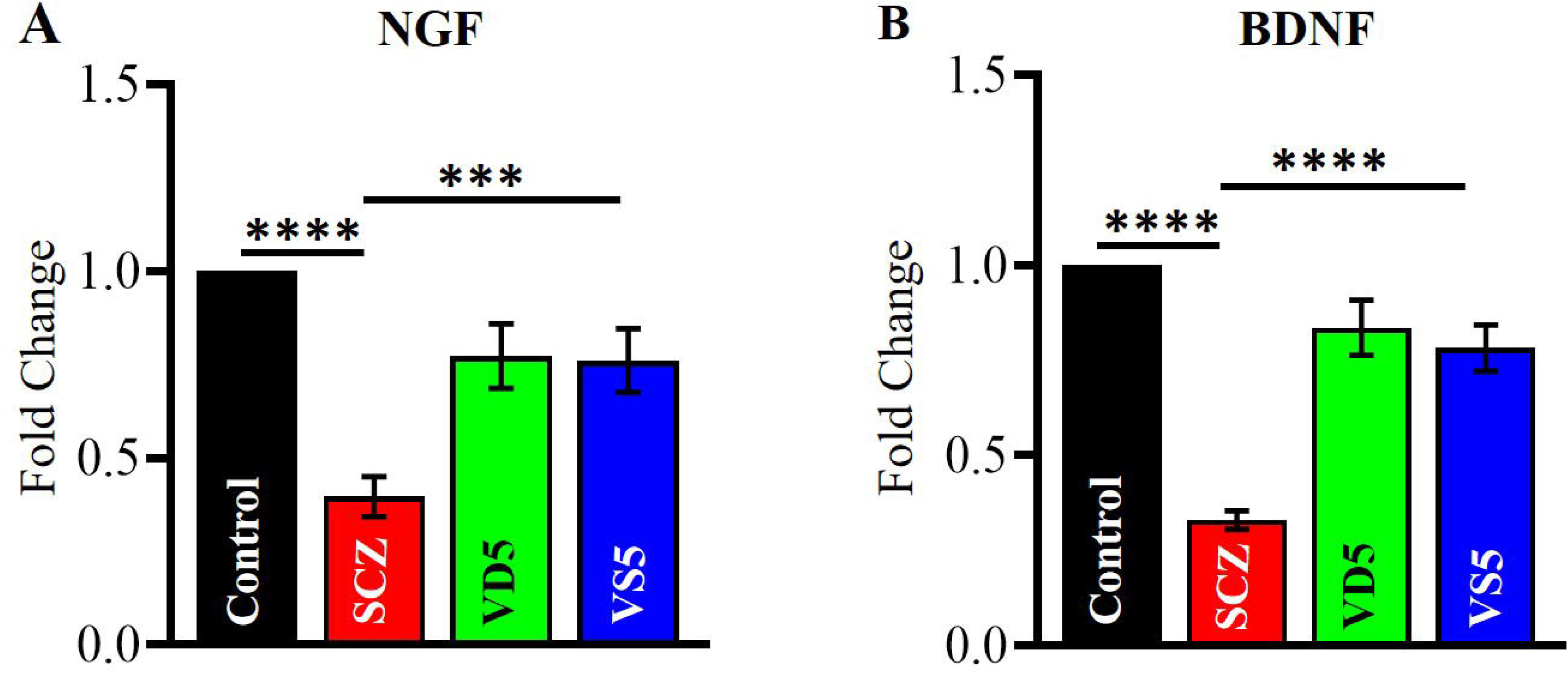
VD intervention rescues neurotrophins, NGF, and BDNF gene expression in SCZ. **(A)** RT-PCR results depicted a significant decrease in the mRNA expression of NGF (Control vs SCZ; n=4, p<0.0001, Student’s pairwise t-test), which was restored upon VD pre-supplementation (SCZ vs VS5; n=4, p<0.001, Student’s pairwise t-test). **(B)** The mRNA expression of BDNF was also observed to be decreased in SCZ mice (Control vs SCZ; n=4, p<0.0001, Student’s pairwise t-test). On VD pre-supplementation BDNF mRNA expression got restored in VS5 mice (SCZ vs VS5; n=4, p<0.0001, Student’s pairwise t-test). Data is normalized, and represented as mean ± SEM.

**Fig. 12.**
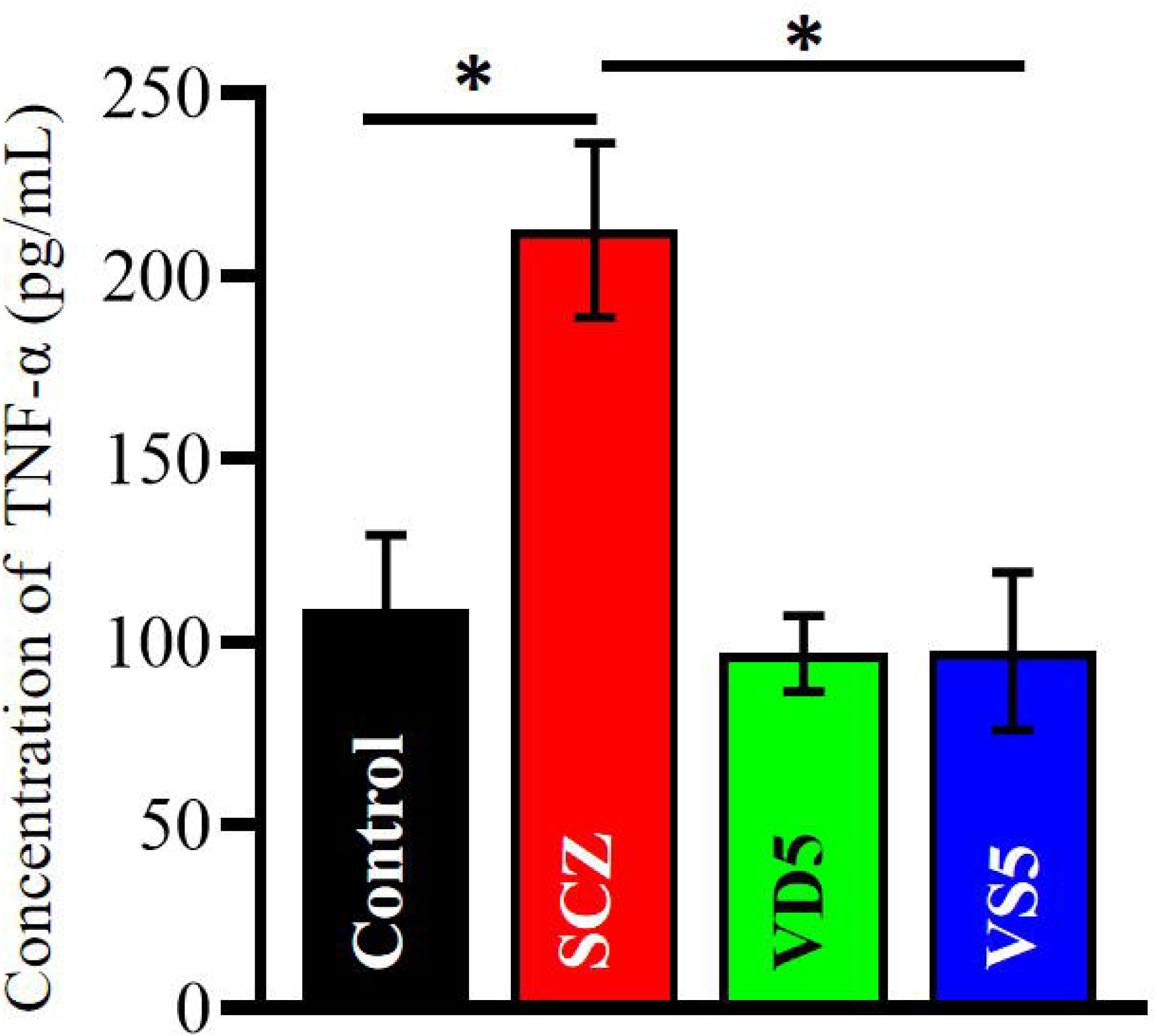
An Anti-inflammatory effect of VD in the PFC of SCZ mice. On the 15^th^ day, MK-801 treatment markedly elevated TNF-α levels in SCZ mice as compared to control (Control vs SCZ; n=5, p=0.02, Students’ pairwise t-test). Conversely, VD pre-supplementation significantly attenuated this increase in VS5 mice when compared to SCZ mice (SCZ vs VS5; n=5, p=0.05, Students’ pairwise t-test). Data represented as mean ± SEM.

### Statistical Analysis

The experimental data are presented as normalized values with respect to the control. The results are represented as bar graphs depicting mean ± SEM for each group of mice. Statistical analysis was performed using two-way repeated-measures ANOVA with power analysis or Student’s pairwise t-test followed by Holm-Bonferroni correction for multiple comparisons wherever applicable using either “R” studio software or GraphPad Prism version 8. A p value of < 0.05 was considered statistically significant (*p <0.05, **p <0.01, ***p <0.001, ****p <0.0001). All the data were graphically represented using GraphPad Prism version 8.

## Results

### Pre-supplementation of Vitamin D3 alleviates hyperlocomotion in SCZ mice

To explore the effect of VD pre-supplementation on anxiety levels that mimics the positive symptoms of SCZ, two-way repeated measures ANOVA with power analysis was performed among the four different groups namely, **Control, SCZ, VD5 and VS5**. A highly significant difference was observed in the total distance travelled by the four different groups of mice (Fig. 2). A prominent group effect was observed (F=54.48, p<0.001) with a power value of 1.00, and a day’s effect (F=75.00, p<0.001) with power value of 0.99. On the 15^th^ day, VS5 mice induced with MK-801, and pre-supplemented with VD showed a significant decrease in the hyper locomotor activity. Overall, two-way repeated measures ANOVA revealed a significant difference in the mean among all the four groups with strong interaction between the groups and day (f _(6)_ =23.18, p<0.001), indicating VD’s beneficial effect in reducing the anxiety in SCZ mice. Similar effects were observed with the other two doses of VD, namely 1000IU/kg and 2000IU/kg (Fig. S2A, S2B). As all the three doses of VD (500IU/kg/day, 1000IU/kg/day, and 2000IU/kg/day) produced a comparable reduction in the locomotor phenotype, we selected the 500IU/kg/day dose for subsequent behavioral and biochemical studies.

### D3 nutraceutical enhances cognitive impairment in MK-801 induced mice

The NOR test showed significant differences in the recognition memory amongst the groups, as measured by the discrimination index (Fig. 3). During the training/exploratory phase, when there was no saline or MK-801 being administered, the respective groups of mice showed no difference in the time spent on the exploration of two similar objects. This indicated that there was no preference for a certain position of the objects in the area. During the recognition session, i.e. the time used by each of group of mice to explore the novel object, SCZ mice took much longer to explore the novel object (Control vs SCZ; 1.00 ± 0.05 vs −0.37 ± 0.11, n=12-14, p=0.0005). This suggests that SCZ mice failed to discriminate the novel from the familiar object, and spent almost equal or less time with the novel object, and demonstrated a severe impairment in the recognition memory. In contrast, control mice showed a strong preference for the novel object and reflected an intact memory function. VD5 mice (supplemented with only 500IU/kg/day of VD) did not significantly differ from the controls group (Control vs VD5; 1.00 ± 0.05 vs 0.63 ± 0.09, n=12-24, p=0.48) and displayed similar preserved recognition memory. Notably, VS5 mice that received VD pretreatment showed a significant rescue in the recognition memory as compared to MK-801 treated mice (SCZ vs VS5; −0.37 ± 0.11 vs 1.04 ± 0.08, n=12-14, p=0.015). VD pre-supplementation to VS5 mice facilitated a robust rescue in the cognitive memory performance, and a strong preference for novel objects as compared to SCZ mice.

### Pre-supplementation of Vitamin D3 rescues memory deficits in SCZ mice

A dysfunctional dorsolateral prefrontal cortex and an impairment in the working memory function are core features observed in schizophrenia (SCZ) patients [88, 89]. To test the efficacy of Vitamin D3 (VD) on memory deficits induced by MK-801 in SCZ mice, we conducted the passive avoidance test (PAT). This test is mainly used to determine the latency (in seconds) i.e., the time taken for mice to enter the dark/shock zone associated compartment during the test phase. The test phase was facilitated 4 hours post training followed with a foot shock. Before the training no significant difference was observed in latency to enter the dark compartment among all the groups of mice (Fig. 4). After 4 hours of training, SCZ mice showed a robust decrease in the latency to enter the dark zone as compared to the control group (Control vs SCZ; 134.25 ± 23.96 vs 53.8 ± 13.11, n=13-17, p=0.028). It reflected that MK-801 caused an impairment in the short-term memory process in SCZ mice. In contrast, VS5 mice showed a significant recovery in the latency when compared to the MK-801 induced SCZ mice, further demonstrating the cognitive enhancer potential of VD in SCZ (SCZ vs VS5; 53.8 ± 13.11 vs 119.64 ± 24.46, n=13-17, p=0.04). Control and VD5 mice exhibited similar retention memory (Control vs VD5; 134.25 ± 23.96 vs 153.76 ± 14.41, n=13-17, p=0.73). These data indicate that nutraceutical interventions like VD can be beneficial especially in neurotoxic conditions that were induced by MK-801 on neurocognitive and memory function [90].

### Modulatory effect of VD on the gene expression of NMDARs and α7nAChRs subunits

An alteration in NMDRs and α7nAChRs are documented to be altered in various pharmacological and genetic rodent models of SCZ [25, 40, 92–101]. These receptors downregulation in expression and function are known contributors to synapse pathology and increased synaptic pruning [102]. SCZ mice showed severe downregulation in the gene expression of NRI subunit of NMDARs (Control vs SCZ; 1.00 ± 0.00 vs 0.573 ± 0.05, n=6, p<0.0001, Fig. 5A). The mRNA expression of NR1 in VS5 group significantly increased upon VD administration as compared to SCZ mice (SCZ vs VS5; 0.573 ± 0.00 vs 1.04 ± 0.15, n=6, p=0.005). Similarly, the NR2A subunit of NMDARs mRNA expression was attenuated in SCZ mice when compared with control mice (Control vs SCZ; 1.00 ± 0.00 vs 0.57 ± 0.06, n=6, p=0.003, Fig. 5B), and got normalized to that of the control mice on pre-supplementation of VD for 15 days (SCZ vs VS5; 0.57 ± 0.06 vs 0.77 ± 0.07, n=6, p<0.0001). NR2B-containing NMDARs was downregulated in SCZ mice (Fig. 5C), and got significantly improved on VD intervention (Control vs SCZ; 1.00 ± 0.00 vs 0.42 ± 0.09, n=6, p<0.0001; SCZ vs VS5; 0.42 ± 0.09 vs 0.69 ± 0.11, n=6, p=0.02).

MK-801 induced SCZ-like mice showed a major reduction in the gene expression of α7nAChRs compared to the control mice (Control vs SCZ; 1.00 ± 0.00 vs 0.44 ± 0.07, n=4, p=0.0002, Fig. 5D), while the pre-supplementation of VD continuously for 15 days substantially rescued the gene expression of α7nAChRs in the PFC (SCZ vs VS5; 0.44 ± 0.07 vs 0.76 ± 0.07, n=4, p=0.02). This normalization of cholinergic receptors gene expression indicates a channel protective role of VD towards the maintenance of synaptic signaling integrity in the PFC. Importantly, no significant changes in the mRNA expression of ionotropic receptors were observed between the control and VD5 supplemented group, indicating that VD administration does not affect receptor alteration under normal physiological conditions.

### VD pretreatment restores α7nAChRs expression in the PFC

Immunohistochemical analysis of PFC showed significant differences in the number of α7 nicotinic acetylcholine receptors (α7nAChRs) positive cells based on the group (Fig. 6). One - way ANOVA analysis revealed a marked reduction in the number of α7nAChRs positive cells in the PFC of SCZ mice when compared to age matched controls (Control vs SCZ; 69.58 ± 5.64 vs 27.65 ± 3.88, n=3 mice, F (3, 31) = 23.4, p<0.0001), characterized by diminished staining intensity and altered distribution across the various cortical layers that are critical for various cognitive functions impaired in SCZ. These findings align with earlier reports of decreased α7nAChRs expression in SCZ and related cognitive issues [105]. VD pretreatment significantly improved the SCZ-induced reduction in the α7 positive cells (SCZ vs VS5; 27.65 ± 3.88 vs 67.09 ± 4.63, n=3 mice, F (3, 31) = 23.4, p < 0.0001) highlighting VD’s neuroprotective role in counteracting cholinergic deficits amid neuroinflammation and oxidative stress, consistent with prior reports. On the other hand, the expression levels of α7nAChRs was indistinguishable from the controls when compared with VD5 mice (Control vs VD; 69.58 ± 5.64 vs 72.11 ± 6.56, n=3 mice, F (3, 31) = 23.4, p<0.0001). A significant change in the α7nAChRs expression however, was observed between the SCZ, and VD5 groups of mice (SCZ vs VD; 27.65 ± 3.88 vs 72.11 ± 6.56, n=3 mice, F (3, 31) = 23.4, p<0.0001). This supports the idea that VD might possibly affect nicotinic acetylcholine receptor expression, and cholinergic signaling in SCZ [106, 107]. Thus, VD sufficiency may represent an effective therapeutic strategy for enhancing cholinergic signaling and α7nAChRs expression as demonstrated in several animal models of Parkinson’s disease (PD), Huntington’s disease (HD), and Alzheimer’s disease (AD) [3].

### VD mitigates AChE activity levels in the PFC of SCZ mice

To assess the effect of VD on cholinergic signaling and neurotransmission, acetylcholinesterase (AChE) activity assay was performed on the pre-frontal cortex samples across the four experimental groups of mice. SCZ mice showed a significant rise in the AChE activity when compared to control mice (Control vs SCZ; 154.9 ± 20.18 mU/mg vs 528.3 ± 101.7 mU/mg, n=5, p=0.01), indicating that MK-801 severely disrupts cortical neurotransmission, consistent with SCZ like pathology. However, administration of VD alleviated MK-801 induced alterations and reduced the AChE activity in SCZ mice (SCZ vs VS5; 528.3 ± 101.7 mU/mg vs 237.3 ± 33.00 mU/mg, n=5, p=0.04). VD pre-supplementation alone did not significantly alter the AChE activity relative to the controls (Control vs VD5; 154.9 ± 20.18 mU/mg vs 204.5 ± 64.55 mU/mg, n=5, p=0.266). These results suggest that VD’s sufficiency exerts an anti-anticholinesterase effect in SCZ (Fig.7). Consistent with our previous observations and by others, VD intervention might help in reducing deficits in cholinergic neurotransmission by decreasing AChE activity and thereby restoration in the ACh levels in SCZ [108].

### A reduction in circulating 25-hydroxyVitamin D3 levels in SCZ

The circulating levels of 25-hydroxyvitamin D3 (calcidiol) were found to be lower in SCZ mice as compared with Controls (n=5). Similarly, the serum calcidiol levels were analyzed in patients with SCZ (n=113) and healthy controls (n=73). Descriptive statistics revealed lower mean calcidiol levels among patients with SCZ when compared to controls. The data revealed a statistically significant difference in the calcidiol levels between both the groups (Control vs SCZ; 21.48 ± 0.00 vs 19.23 ± 0.00, U=3299.5; p=0.021, Mann-Whitney U test). The rank-biserial correlation was calculated to be 0.20, indicative of a small-to-moderate effect size, suggesting that VD levels tend to be lower in individuals with SCZ though the magnitude of difference could be modest (Fig. 8). These findings support previous evidence implicating VD insufficiency in SCZ and emphasize the need to consider VD status in clinical management, and research on psychosis [17, 109].

### Calcitriol pre-treatment enhances alpha7 nicotinic acetylcholine single-channel currents

In order to explore whether VD may directly modulate ion-channel function, α7-Venus tagged nicotinic acetylcholine receptors was transfected in HEK293T cells (see methods) with 1μM calcitriol for 24h. Cells were patched in cell-attached configuration with 1mM nicotine in the recording pipette. 4 GΩ seal and above cells were included only for recording (Fig. 9).

The single-channel recordings in cell-attached configuration from control cells, and 1μM calcitriol treated cells were performed on second- and third-day post-transfection. The single-channel α7 receptors mediated currents were activated by nicotine (1mM) present in the patch pipette electrode solution. The extracellular solution was identical to the patch pipette recording solution except the presence of nicotine. The patch of membrane in cell-attached mode was voltage-clamped at a pipette potential of +40 mV. The estimated transmembrane potential was around 100 mV as documented previously [84]. A substantial enhancement in the normalized mean peak current amplitude of α7nAChRs was observed from 1μM calcitriol pretreated HEK293T cells as compared to that of control (Control vs calcitriol; 1.00 ± 0.15 vs 1.73 ± 0.21, n=4, p=0.02). Our results suggest a direct effect of calcitriol the functionality of α7nAChRs. These results can further be speculated that enhanced functionality of α7nAChRs in the PFC may enhance synaptic efficacy and neurotransmission via its interaction with NMDARs by a cross-talk mechanism as proposed in earlier studies [93, 110–113]. Targeting α7nAChRs may alleviate NMDA receptor hypofunction and rescue synaptic functions as observed in SCZ [92]. Vitamin D3 sufficiency in SCZ may be proposed to contribute towards the maintenance of a given pool of functional α7nAChRs receptors that may help to alleviate the different psychosis symptoms.

### D3 nutraceutical restores PSD-95 gene expression in MK-801 induced mice

Postsynaptic density protein-95 (PSD-95) constitutes one of the major regulators in stabilizing and trafficking of N-methyl-D-aspartic acid receptors as shown in earlier reports (NMDARs)[114–116]. To explore the synaptic protective effects of VD pre-supplementation in SCZ mice, we analyzed the protein expression of PSD-95, Syntaxin, and SNAP-25 (Fig. 10). PSD-95 protein levels were markedly reduced in the SCZ group as compared to control groups of mice (Control vs SCZ; 1.00 ± 0.00 vs 0.66 ± 0.05, n=4, p<0.0001). Pre-supplementation with VD robustly restored the PSD-95 protein expression in VS5 mice (SCZ vs VS5; 0.66 ± 0.05 vs 1.05 ± 0.149, n=4; p=0.04). However, Syntaxin and SNAP-25 expression showed no significant changes following VD pre-supplementation. Previous study performed using MK-801showed a similar perturbation in the synaptic components in PFC [117]. In addition, to PSD-95, we tested the serine-threonine phosphatase, Pan-calcineurin expression levels in PFC documented to be dysregulated in SCZ [118]. We found Pan-calcineurin levels were significantly elevated in SCZ mice relative to controls (Control vs SCZ; 1.00 ± 0.00 vs 1.48 ± 0.22, n=6, p=0.04) and were rescued back upon VD pre-supplementation (SCZ vs VS5; 1.48 ± 0.22 vs 1.10 ± 0.12, n=6, p=0.03). The biological nuclear receptor known to induce genomic signaling of VD i.e. Vitamin D receptors (VDRs) were also reduced in the SCZ group, and increased in VS5 mice (Control vs SCZ; 1.00 ± 0.00 vs 0.82 ± 0.04, n=5, p=0.002; SCZ vs VS5; 0.82 ± 0.04 vs 1.05 ± 0.11, n=5, p=0.03). Our findings support previous reports demonstrating that VD enhances PSD-95 protein expression in the PFC via BDNF/TrkB-mediated protein synthesis [66]. Since, calcineurin is known regulator of synaptic elements [119], our data suggest VD as a mediator for a large synaptic interactome complex and can restore impaired synaptic communication in SCZ (Fig. 10).

### Neurotrophins enhancer effect of VD in SCZ mice

Previous work from our lab has documented a neurotrophic enhancer effect of Vitamin D3 (VD) post-supplementation in Huntington’s disease (HD) mouse model [3, 45–47]. Recent clinical study documented a significant reduction in the serum levels of two major neurotrophins namely, brain-derived neurotrophic factor (BDNF), and nerve-growth factor (NGF) in male patients with chronic schizophrenia when compared to the healthy controls [120]. SCZ mice displayed a significant reduction in the gene expression of NGF when compared to control mice (Control vs SCZ; 1.00 ± 0.00 vs 0.39 ± 0.05, n=4, p<0.0001). VD pre-supplementation robustly rescued NGF gene expression in VS5 mice relative to SCZ mice (SCZ vs VS5; 0.39 ± 0.05 vs 0.76 ± 0.08, n=4, p<0.001), indicating that MK-801 induction did not compromise VD’s biological effects. Notably, VD treatment alone did not significantly alter the NGF expression when compared with control (Fig. 11A).

The mRNA expression of BDNF also exhibited a comparable pattern of dysregulation (Fig. 11B). SCZ mice showed a significant decrease in the gene expression as compared to control mice (Control vs SCZ; 1.00 ± 0.00 vs 0.33 ± 0.02, n=4, p<0.0001). VS5 mice that received VD pre-supplementation for 15 days were rescued its expression (SCZ vs VS5; 0.33 ± 0.02 vs 0.78 ± 0.06 vs, n=4, p<0.0001). Collectively, our data indicates that VD pre or post supplementation has a neurotrophins enhancer effect in different neuropathological conditions [45, 120] and it parallels another finding by Khairy and colleagues who demonstrated a restoration in the gene expression of BDNF in an aged rodent model [121].

### Identification of Putative Vitamin D Response Elements (VDREs) in target gene promoters

*In-silico* analysis of the promoter regions of the selected genes using the JASPAR database identified multiple putative Vitamin D Response Elements (VDREs). The binding motifs corresponding to VDR and/or the VDR: RXR heterodimer were detected within the promoter regions of various NMDA receptor subunits namely, NR1, NR2A, and NR2B. Other targets included α7 nicotinic acetylcholine receptors (α7nAChRs), nerve growth factor (NGF), brain-derived neurotrophic factor (BDNF), post-synaptic density protein-95 (PSD-95), and inflammatory cytokine, tumor necrosis factor alpha (TNF-α). Several high scoring binding sites at 75-80% relative profile score threshold were identified, suggesting potential transcriptional regulation of these genes by Vitamin D3 signaling. The predicted VDREs were distributed at different positions relative to the transcription start site (TSS) and were present on both the positive and the negative strands. Detailed information regarding the motif locations, strand orientation, and the relative profile scores is presented in Table. 3.

### Anti-inflammatory potential of “D3” nutraceutical in SCZ

SCZ mice injected with MK-801 showed a profound enhancement in the levels of TNF-α as compared to control mice (Control vs SCZ; 108.9 ± 20.28 vs 212.5 ± 23.82 pg/mL; n=5, p=0.02, Fig. 12). Pre-supplementation of VD significantly reduced the TNF-α levels in SCZ mice (SCZ vs VS5; 212.5 ± 23.82 pg/mL vs 97.45 ± 21.58 pg/mL; n=5, p=0.02) while VD supplementation alone did not significantly alter the TNF-α levels relative to the controls (Control vs VD5; 108.9 ± 20.28 vs 96.88 ± 10.33, n=5, p=0.98) (Fig. 12). These results suggest the anti-inflammatory potential of VD in myriad neuropathological conditions [45, 68, 123, 124].

## Discussion

The World Health Organization predicts that neuropsychiatric illnesses like schizophrenia (SCZ) and other age-related mental disorders are going to rise substantially in the next two decades [125]. SCZ is a multifactorial mental health disorder and many symptoms arise due to N-methyl-D-aspartate receptor (NMDAR) hypofunction [97, 126, 127]. Along with glutamatergic abnormalities, cholinergic dysfunction and loss of alpha7 nicotinic acetylcholine receptors (α7nAChRs) are also observed in SCZ [94, 95]. Hence, the diagnosis and identification of molecular targets that can restore these neurotransmitter receptors function and expression appears to be at the forefront of brain research [128]. These excitatory ionotropic receptors constitute an important element of excitatory synapses, and facilitates neuronal communication and synaptic plasticity [111, 116, 129, 130]. Their downregulation has been associated with numerous neurological and psychiatric disorders including SCZ [34, 115, 116, 128, 131].

Hypovitaminosis of “D3” nutraceutical appears to be associated with a number of neurodegenerative and neuropsychiatric disorders like Alzheimer’s disease (AD), Parkinson’s disease (PD), Huntington’s disease (HD), schizophrenia (SCZ), depression, mood disorders, attention-deficit hyperactivity disorders, and many others [56, 132–137]. The present work elucidates a cheap nutraceutical intervention, Vitamin D3 (or cholecalciferol) pre-supplementation, as a strategy towards the alleviation of different subunits of N-methyl-D-aspartate receptors (NMDARs), αlpha7 nicotinic acetylcholine receptors (α7nAChRs), post-synaptic density protein-95 (PSD-95), and various behavioral deficits induced by MK-801 causing NMDARs hypofunction in a mouse model of SCZ [138, 139]. “MK-801” is a known NMDA receptor non-competitive antagonist, and widely used as a psychomimetic drug to recapitulate motor syndrome, hyperlocomotion, and cognitive impairments in rodent models [39–41, 138, 140]. We utilized MK-801 to induce synaptic, neurotrophic and ionotropic receptor changes as reported previously by various studies [30, 41, 139, 141].

Cognitive deficits are one of the core features of schizophrenia (SCZ) [38]. The current antipsychotic medications have shown limited efficacy on cognitive impairments [86]. SCZ has been consistently associated with elevated tumor necrosis factor- α (TNF-α) levels, highlighting the contribution of immune-inflammatory pathways to its pathophysiology [122]. Our findings depicted that SCZ mice presupplemented with 500IU/kg/day of VD (i.e. VS5 mice) displayed decreased anxiety, and enhanced cognitive ability (Fig. 2, 3 and 4) due to a restoration in the gene expression of major N-methyl-D-aspartate receptor subunits (NMDARs), namely, NR1, NR2A, and NR2B subunits Fig. 5A, 5B and 5C). Our results are consistent with the *in-vivo* findings by another group where a similar dose of VD (500IU/kg/day) improved cognitive function, and cholinergic transmission in streptozotocin-induced diabetic rats [42]. It parallels recent reports that showed the neuroprotective effects of VD supplementation in monosodium glutamate induced behavioral, cholinergic, oxidative and neuroinflammatory deficits in rats [85]. VS5 mice exhibited a rescue in the acetylcholine neurotransmitter levels together with increased expression of αlpha7 nicotinic acetylcholine receptors in the PFC (α7nAChRs, Fig. 5D, 6, and 7). The restoration of NMDARs subunits together with α7nAChRs may compensate for the hypofunction circuitry of PFC, and downregulation of these receptors probably contributed to all the above behavioral deficits as observed in MK-801 induced SCZ-like mice [96, 97, 104]. A direct effect of calcitriol pretreatment to HEK293T cells transfected using alpha-7-Venus tagged fluorophore displayed a robust increase in the peak single channel current amplitude (Fig. 9). The enhanced expression and function of this cholinergic receptor suggests that VD rescues synaptic communication through ionotropic receptors in neuropathological conditions like SCZ. Its sufficiency can enhance synaptic connection by restoration of glutamatergic signaling via NMDARs subunits, PSD-95, and α7nAChRs that is likely to form a supramolecular complex in the PFC circuitry (Fig. 13) [27, 142]. The importance of α7nAChRs comes from extensive studies that documented α7nAChRs-NMDARs cross-talk or where ligand induced activation of this cholinergic receptor have potentiated the glutamatergic currents [93, 101, 111]. A decrease in the α7nAChRs expression is known to contribute to gating deficits observed in SCZ [92, 95, 103]. Since α7nAChRs undergo significant downregulation in SCZ, causing cognitive deficits and other symptoms, a multidrug-conjugated approach utilizing VD and deeper insights into α7 structure-function analysis may lead to better remedies in the field of SCZ and other neuropsychiatric disorders [92].

**Fig. 13.**
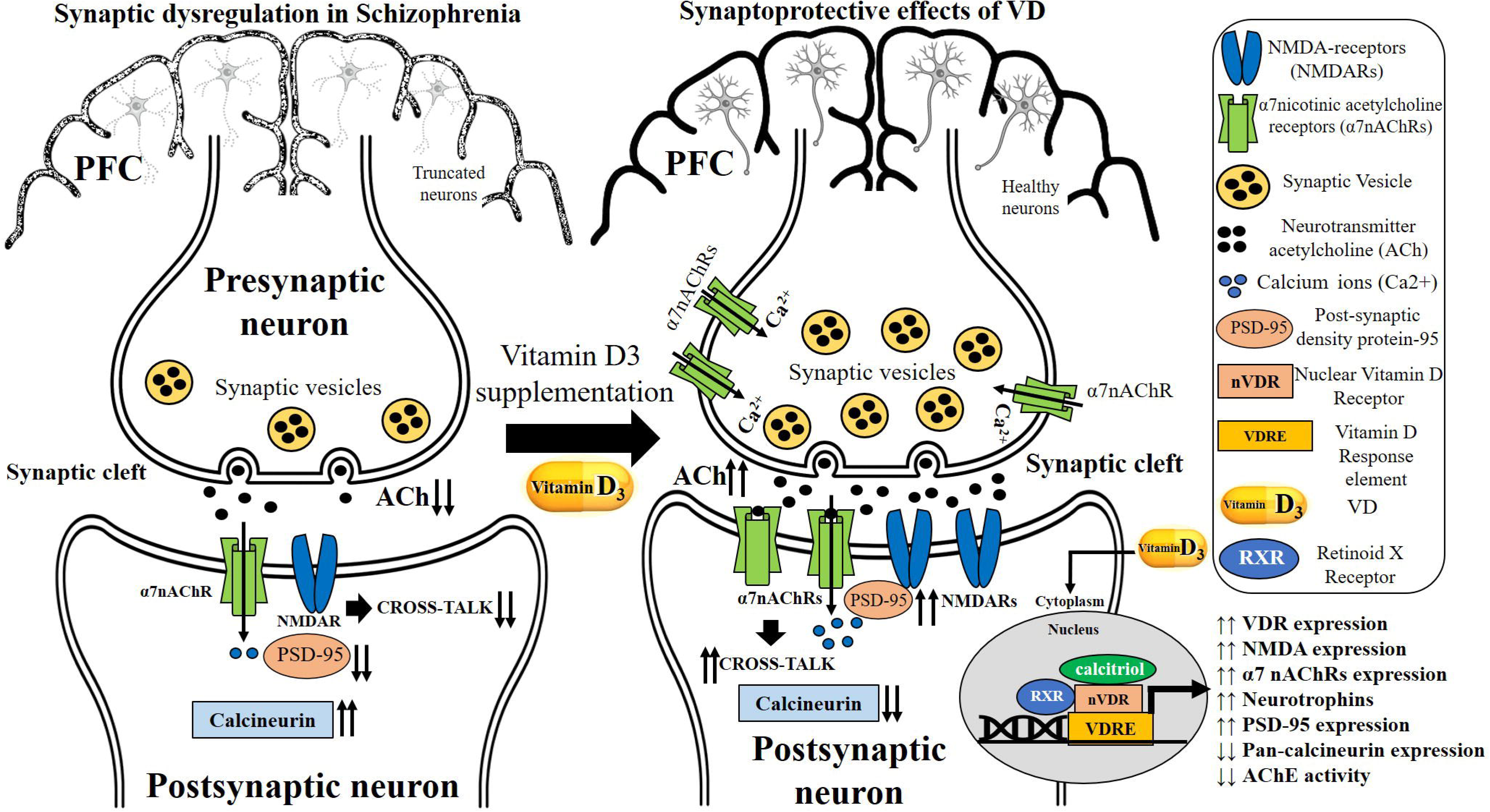
Diagrammatic representation of synaptic enhancement mediated by VD via enhanced α7nAChRs and NMDARs expression. The left panel illustrates synaptic dysregulation associated with schizophrenia (SCZ), characterized by neuronal atrophy, reduced expression of α7 nicotinic acetylcholine receptors (α7nAChRs), impaired N-methyl-D-aspartate receptor (NMDA) signalling, reduced levels of postsynaptic density protein-95 (PSD-95), and disrupted calcium homeostasis within the prefrontal cortex (PFC). Collectively, these alterations contribute to compromised cholinergic neurotransmission, impaired synaptic plasticity and deficits in neuronal communication. The right panel depicts the proposed neuroprotective and synaptoprotective effects of Vitamin D3 (VD) pre-supplementation in SCZ. The genomic effects of VD are represented when its active metabolite, calcitriol VD binds to its nuclear Vitamin D receptor (VDR), heterodimerizes with the retinoid X receptor and binds to Vitamin D response elements (VDREs). These bindings further regulate increased transcription of various genes involved in neuronal survival, neuroplasticity, neurotransmission and synaptic maintenance. The activation of VDR-dependent signalling cascade enhances the expression of NMDARs, α7nAChRs, neurotrophic factors (NGF, BDNF), and PSD-95 leading to improved receptor function, enhanced synaptic architecture and stabilisation of postsynaptic complexes. The restoration of α7nAChRs in pre and postsynaptic neuron may then facilitate rescue in neuronal communication in SCZ. In parallel, increased α7nAChRs-NMDARs crosstalk with a downregulation in pan-calcineurin protein expression, and acetylcholinesterase (AChE) activity represents a comprehensive framework through which VD may mitigate SCZ associated synaptic abnormalities and promote restoration of neuronal connectivity and synaptic plasticity in the PFC. Upward arrows indicate increased expression or activity, whereas downward arrows indicate reduced expression or activity. Abbreviations - AChE, Acetylcholinesterase; α7nAChR, α7 nicotinic acetylcholine receptor; BDNF, Brain-derived neurotrophic factor; NGF, Nerve growth factor; NMDA, N-methyl-D-aspartate; NMDAR, N-methyl-D-aspartate receptor; PFC, Prefrontal cortex; PSD-95, Postsynaptic density protein-95; RXR, Retinoid X receptor; SCZ, Schizophrenia; VD, Vitamin D3; VDR, Vitamin D receptor; VDRE, Vitamin D response element.

Synaptic integrity is dependent on the proper function and expression of NMDA receptors, α7nAChRs, and other ion-channels at the synapses, proposed to be dysregulated in SCZ [33, 93]. NMDA receptors are known to function as heteromeric receptors with an essential NR1 subunit, and its deletion causes psychosis-like behaviors and memory dysfunction [98, 127]. It will be worthwhile to test the efficacy of this nutraceutical in transgenic murine models of SCZ, too. A sufficient concentration of calcidiol in the CNS has the ability to preserve the synaptic availability of ionotropic receptors and other synaptic proteins in SCZ. This hypothesis is entirely consistent with the findings of many groups who argued on the efficacy of optimal VD status towards vital brain functions, and aging [14, 58, 143–147]. Thus, VD deficiency can alter brain development and has been projected as a ‘‘sufficient cause’’ with respect to the development of neuropsychiatric disorders such as schizophrenia [11, 18].

### Limitations

Though we provide a promising effect of VD pre-supplementation in the MK-801 induced mouse model of SCZ, our findings have some limitations. It is vital to understand that multiple pieces of evidence pinpoint a time and dose-dependent effects of MK-801 in rodent models [40, 138]. Hence, the efficacy of VD on psychosis symptoms generated by this non-competitive inhibitor of NMDA in PFC may vary accordingly. We explored the pre-supplementation effect of VD only for 15 days (0^th^ day to 15^th^ day, Fig. 1) in SCZ mainly because the half-life of calcidiol is about 21 days. The serum levels of calciferols (i.e calcidiol and calcitriol) may well differ from the endogenous synaptic levels of calcitriol as both neurons and glial cells express major VD metabolic pathway enzymes [97]. Hence, the plasma concentration of calcidiol may not truly reflect the synaptic levels of calcitriol that may affect the major excitatory neurotransmitter receptors modulation in the PFC of VS5 mice. Also, the best acceptable level of VD sufficiency is to map calcidiol levels in the blood, whose half-life is projected to be about 21 days [41]. A recent clinical study deciphered that high doses of VD (4000 IU, and 8000 IU) after a 30-day discontinuation could maintain a better average serum concentration of calcidiol (131.5 nmol/l and 144.10 nmol/l) than the lower doses [109]. The upper safe limit of VD proposed for children is 2000 IU /day, and for adults, can be increased to 10,000 IU/day. In patients with severe mental impairments like depression, even higher doses are suggested to be acceptable [110]. Hence, we used 500IU/kg/day of VD in the present neuropharmacological mouse model of SCZ (Fig. S5).

### Conclusion

The psychotomimetic effect of SCZ induced by MK-801 on Vitamin D3 (VD) intervention benefited locomotion, cognition, neurotrophins, synaptic proteins, and glutamate and cholinergic receptor expression in SCZ. The pre-supplementation of VD was sufficient to restore the behavioral abnormalities observed in SCZ mice. VD restored acetylcholine levels and may be projected as a vital dietary intervention that could rescue some psychiatric symptoms in SCZ. Our transcriptional, translational, and electrophysiological data depicts the neuroprotective effect of VD on two major excitatory ligand-gated ion channel expression, implicated to be dysfunctional in SCZ [97, 127, 148]. VD intervention may increase synaptic activity via upregulation of NMDARs and α7nAChRs in the PFC. VD’s synaptic sufficiency may decrease synaptic pruning in the PFC by increasing synaptic connections that may possibly be projected as one of mechanisms of rescue to cognitive deficits observed in our SCZ mice. VD pre-supplementation reduced the SCZ like phenotypes in the present MK-801 induced SCZ mice via an enhancement in cholinergic neurotransmission. This further supports our hypothesis that VD has a neurotrophic, synaptic protein enhancer, and anti-inflammatory potential in SCZ. VD can show a synaptic resilience effect and may pause or rescue the augmented synaptic pruning in this neuropathological condition that needs further investigation in genetic and neurodevelopmental models of SCZ. It may be speculated that VD sufficiency may have restored the synaptic efficacy of PFC circuitry through a supramolecular interactome complex that included PSD-95, NMDARs, and α7nAChRs. PSD-95 which in turn could have normalized the neuronal circuitry in the PFC (Fig. 13) [33, 35, 114, 149, 150]. Furthermore, VD mediated attenuation of calcineurin may have further enhanced NMDARs function in the PFC [151]. This molecular restoration may be argued to be responsible for the alleviation of behavior deficits observed in SCZ mice (Fig. 2, 3, and 4). Furthermore, VD deficiency was explored in the Indian SCZ patients. The beneficial effect of VD consumption in neuropsychiatric disorders like SCZ however, merits additional investigation and follow-up studies in alleviation of the psychosis symptoms in these patients.

## Acknowledgments

We extend deep gratitude to Professor Raad Nashmi, University of Victoria, British Columbia for lending us cDNA for alpha7 Venus fluorophore, and RIC-3. DBT Builder (BT/INF/22/SP42551/2021) is acknowledged for providing mouse and electrophysiology facilities to conduct behavioural tests, brain dissection and single-channel analysis. The contribution of Central Instrumentation Facility (CIF), Birla Institute of Technology and Science-Pilani (BITS-Pilani), Hyderabad Campus, is gratefully acknowledged for providing access to BD 796 FACS Aria II, Leica confocal microscope (Leica SP8/DMi8), RT-PCR system (BIORAD CFX Opus 96). We also acknowledge the DDT-FIST (SR/FST/LS1-526/2012) facility of the Department of Biological Sciences, BITS-Pilani, Hyderabad campus for providing access to RT-PCR (LC480 Roche, Roche), imaging system for 799 Western blots (Fusion Pulse 6, Vilber Lourmat). We are also thankful to the Department of Health Research (DHR), Government of India (GOI), for the funds and facilities at IICT, and MRU-DIMHANS.

## Availability of data and materials

Data will be provided on reasonable request

## Declarations Ethics approval and consent to participate

Not applicable.

## Consent for publication

Not applicable.

## Author Contributions

PK and OE contributed to the conception and design of the study, methodology, literature review, data analysis, interpretation of results, and analyzed the data for most of those experiments. OE performed all the western blot, behavior, biochemical, and gene expression experiments, analyzed all the data, and contributed to writing the section “Materials and Methods” and results for the manuscript. KG performed single-channel recording. SAM conducted some behavior, and RT-PCR data. VAY and RN performed clinical data on SCZ. SC, SP and RK procured immunohistochemistry, and *in-silico* data. PK contributed to funding acquisition, conceived the research questions, data collection, contributed to the design of experiments, analyzed single-channel electrophysiology data, and wrote the final draft of the manuscript. All authors reviewed and approved the manuscript.

---

**Fig. S1. No effect of VD Pre-supplementation on the body weight of mice.** The body weight of mice in all four experimental groups was monitored throughout the study period. No significant differences in body weight were observed among the groups, indicating that neither MK-801 administration nor VD pre-supplementation affected general health or growth.

**Fig. S2A. Pre-supplementation of 1000IU/kg/day of VD reduces hyper locomotor activity in MK-801 induced SCZ mice.** The total distance travelled differed significantly across the four groups: **Control, SCZ, VD1, VS1** (two-way repeated measures of ANOVA, group effect: F = 44.49, p<0.001; day effect: F=93.04, p<0.001). Mice receiving VD at 1000IU/kg exhibited a marked decrease in hyperlocomotion. The significant group by day interaction (F=35.55, p<0.001) suggests that VD effectively mitigated hyperlocomotion in SCZ mice.

**Fig. S2B. Pre-supplementation of 2000IU/kg/day of VD reduces hyper locomotor activity in MK-801 induced SCZ mice.** The total distance travelled showed a significant difference among the four groups: **Control, SCZ, VD2, VS2** (two-way repeated measures of ANOVA, group effect: F=52.58, p<0.001; day effect: F=93.37, p<0.001). The administration of VD at 2000IU/kg significantly lowered hyperlocomotion in SCZ mice. A marked group and day interaction were also found across the experimental groups where VS2 mice demonstrated a significant improvement in hyperlocomotion activity (F=33.42, p<0.001).

**Fig. S3. Video recordings of open field test across all the experimental groups.** Representative videos showing the hyperlocomotor activity of MK-801 induced SCZ mice and its attenuation following VD pre-supplementation.

**Fig. S4. Video recordings of novel object recognition test across all the experimental groups.** Representative videos showing the exploratory behavior across all the experimental groups of mice, highlighting impaired novel object preference in SCZ mice and its improvement following VD pre-supplementation in VS5 mice.

**Fig. S5. VD recommended dose for clinical and preclinical studies.** Comparison of VD doses used in humans and rodents, highlighting recommended intake, supplementation ranges, and on the selection of the VD dose utilized in the present study. The mouse dose of 500 IU/kg/day corresponds to a clinically relevant human-equivalent dose recommended in severe mental health illnesses like SCZ.

**Table 1:**
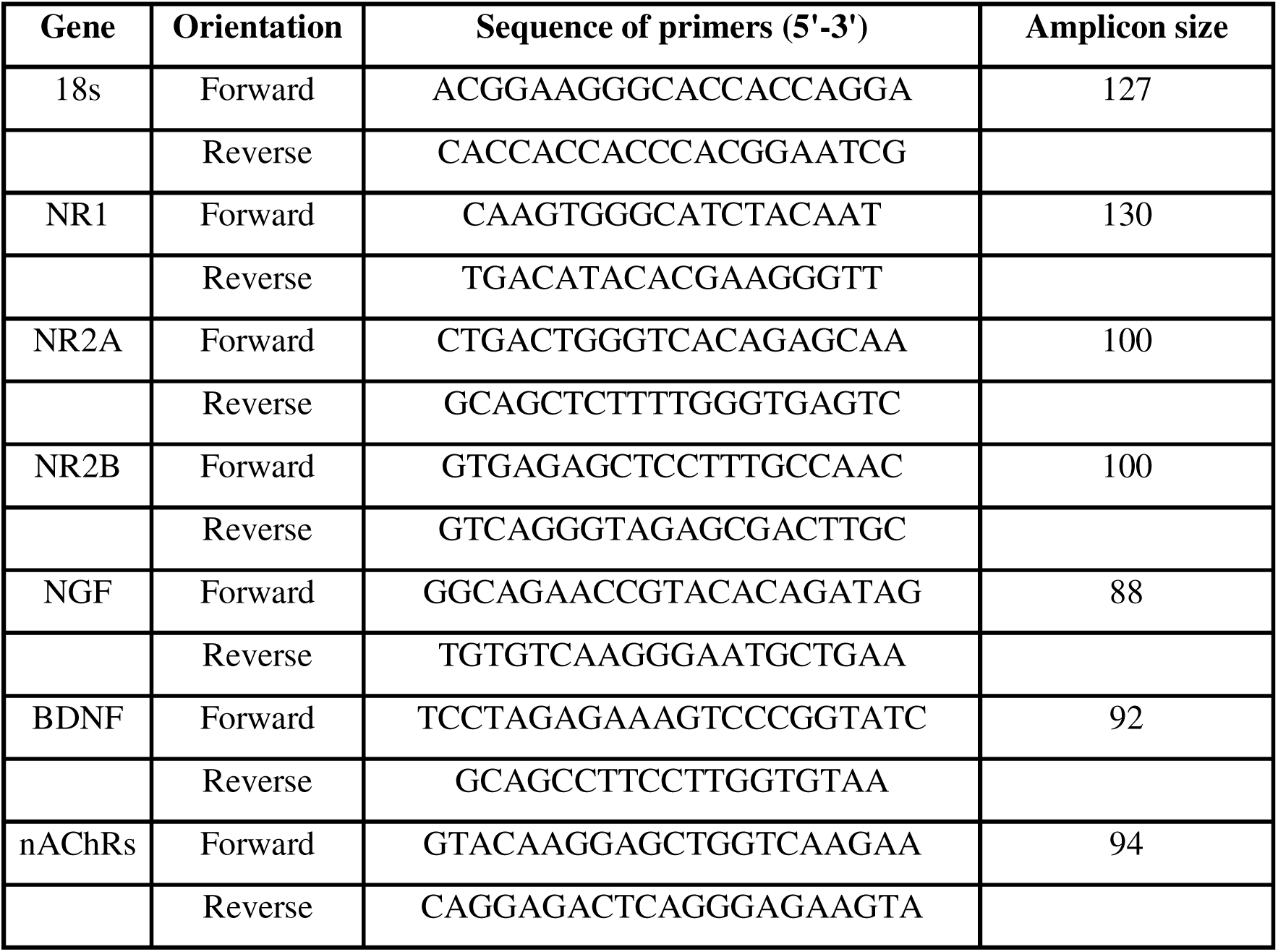
Sequence of primers used in RT-PCR studies.

**Table 2:** List of antibodies used in Western blotting.

| **Antibody** | **Supplier** | **Catalogue number** | **Dilution** |
| --- | --- | --- | --- |
| GAPDH (D16H11) XP Rabbit mAb., Primary | Cell Signaling Technology | 5174 | 1:1000 |
| Vitamin D3 Receptor (D2K6W) Rabbit mAb., Primary | Cell Signaling Technology | 12550 | 1:1000 |
| PSD95 (D27E11) XP Rabbit mAb., Primary | Cell Signaling Technology | 3450 | 1:1000 |
| Syntaxin 1A (D4E2W) Rabbit mAb., Primary | Cell Signaling Technology | 18572 | 1:1000 |
| SNAP 25 (D7B4) Rabbit mAb., Primary | Cell Signaling Technology | 5308 | 1:1000 |
| Pan Calcineurin A Antibody., Primary | Cell Signaling Technology | 2614 | 1:1000 |
| Anti – rabbit IgG, HRP-linked Antibody., Secondary | Cell Signaling Technology | 7074 | 1:5000 |

**Table 3:**
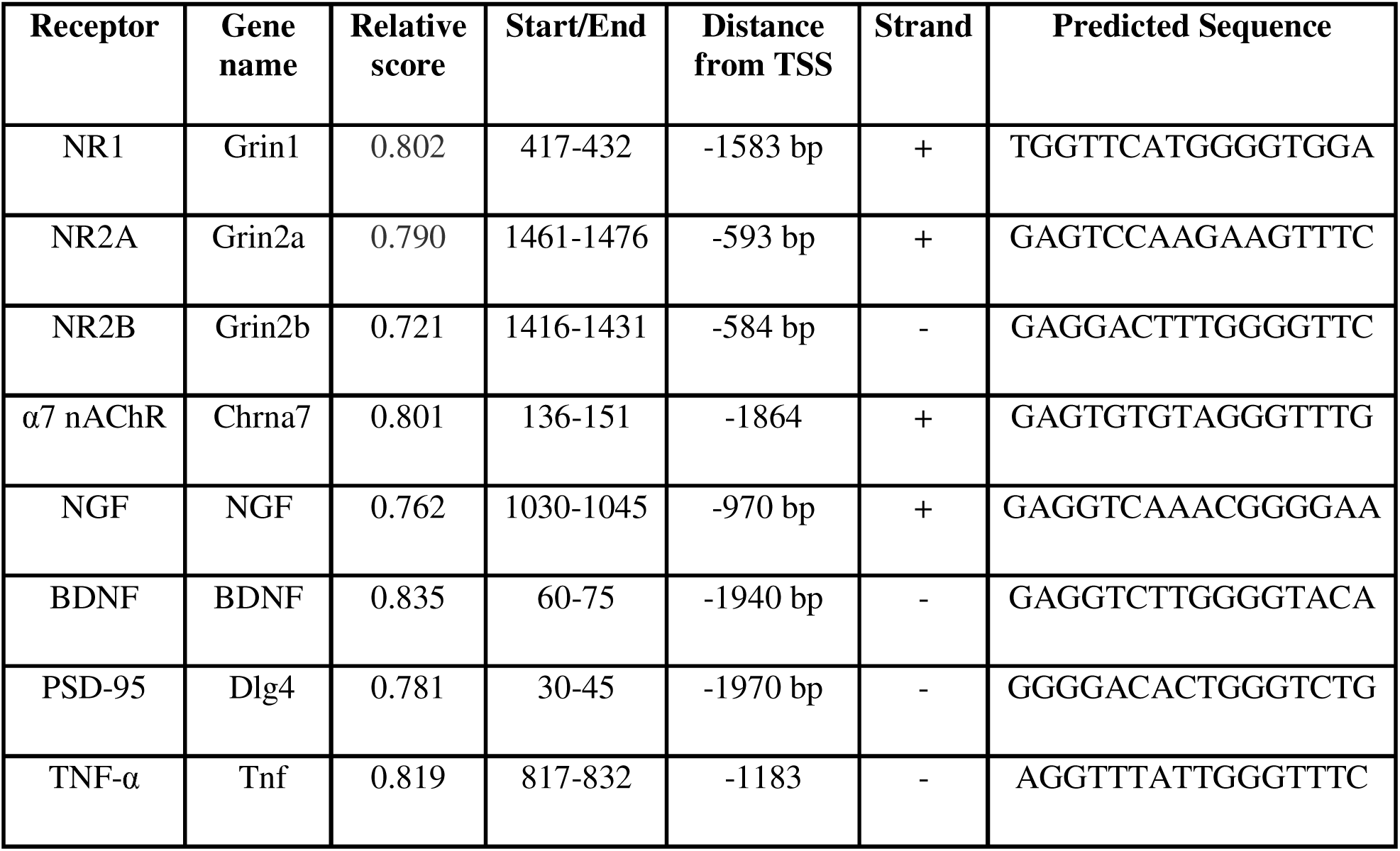
*In-silico* identification of putative Vitamin D Response Elements (VDREs) in the promoter regions of target genes using the JASPAR database.

