## Supplementary figures and images for "D3 nutraceutical pre-supplementation ameliorates behavioral, ionotropic receptors, and synaptic proteins alterations in a MK-801 induced mouse model of schizophrenia"

### Supplemental figure S1

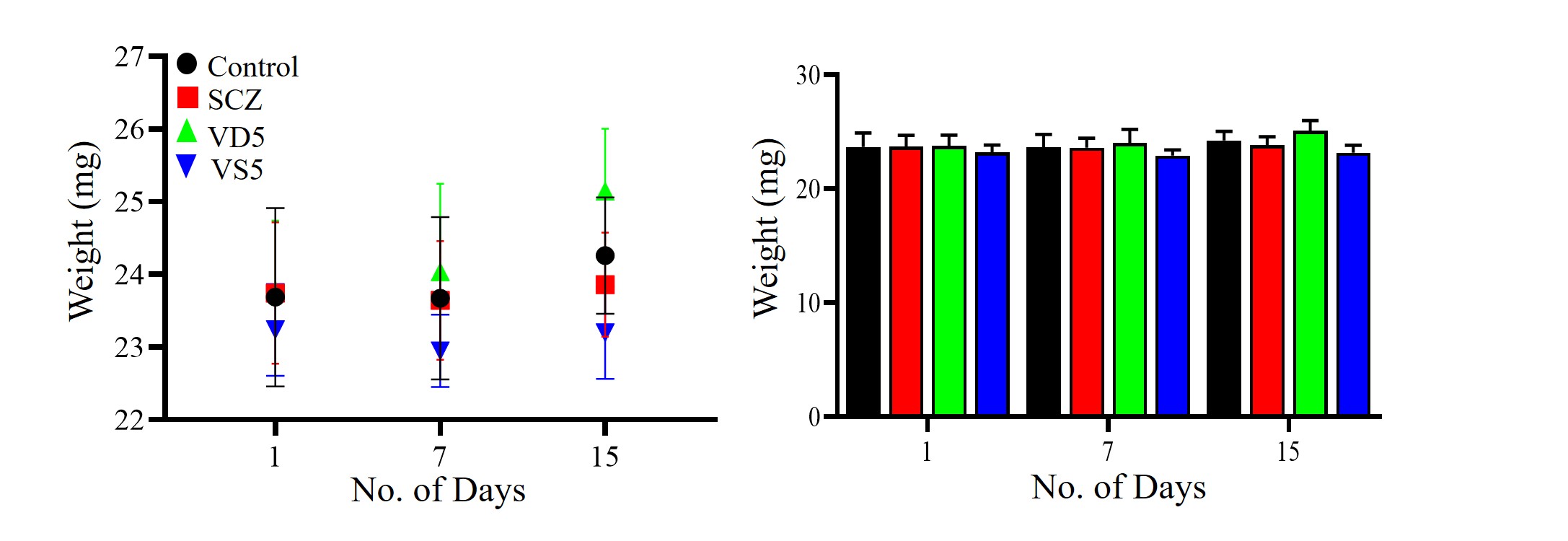

### Supplemental figure S2A

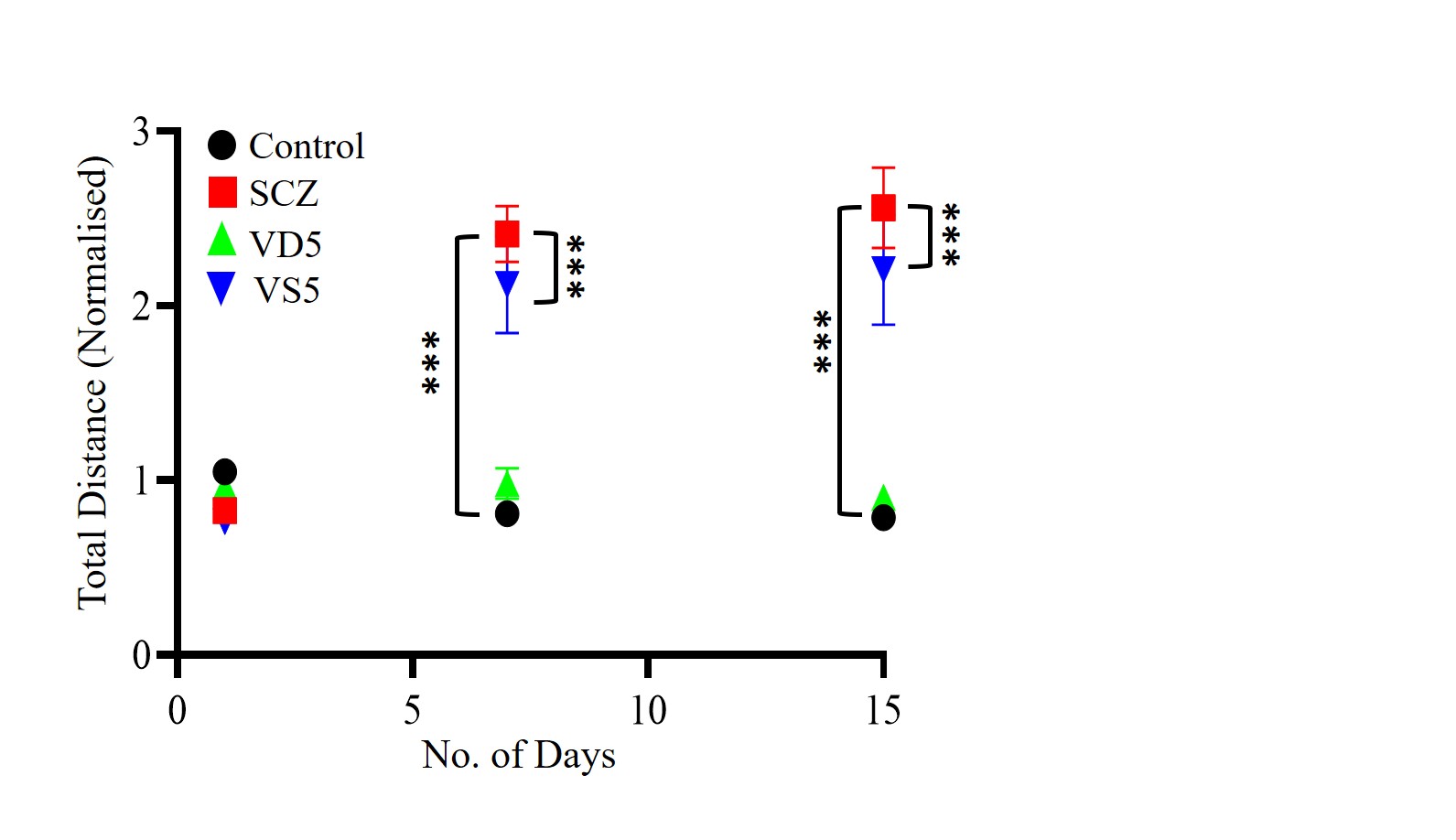

### Supplemental figure S2B

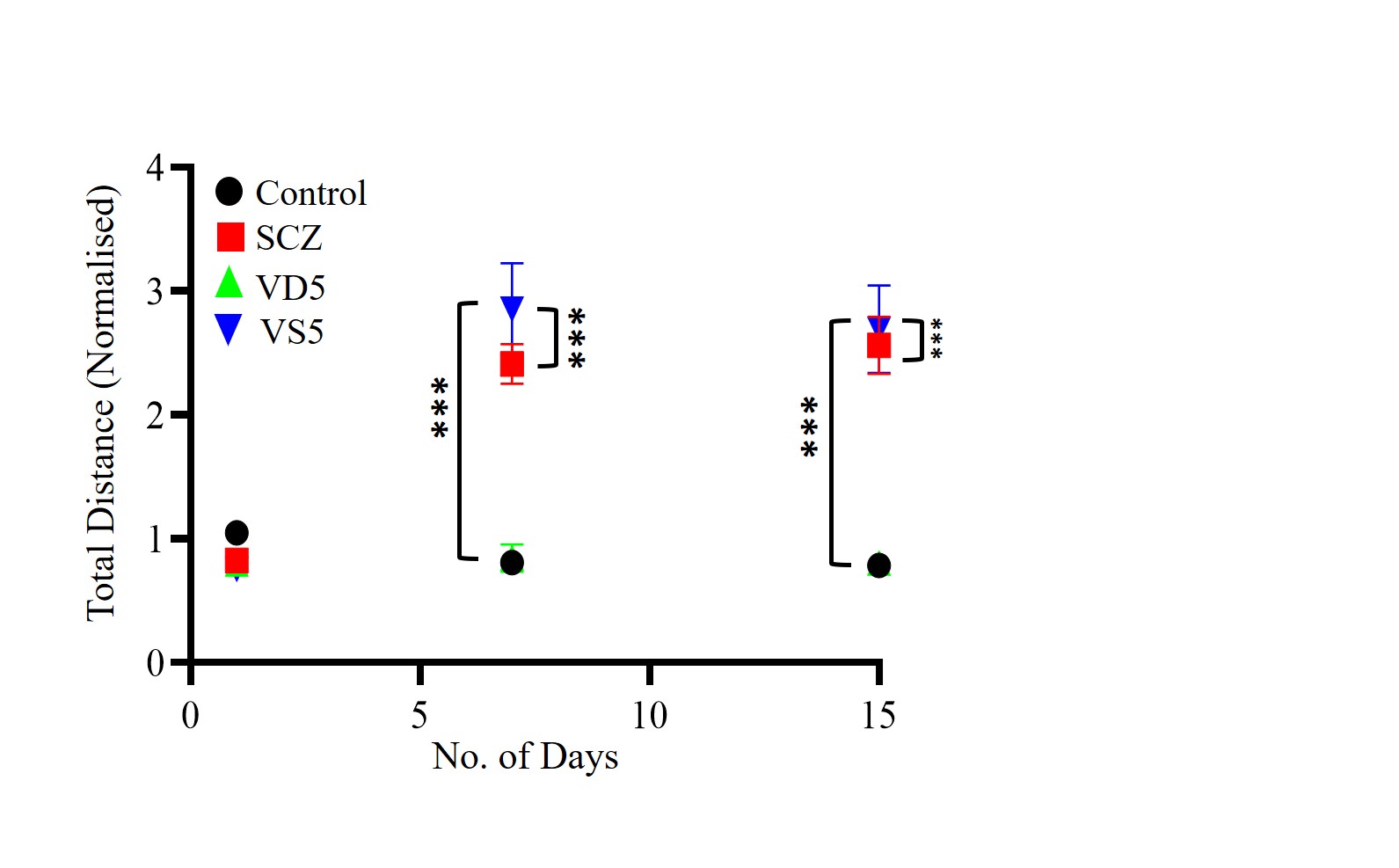

### Supplemental figure S5

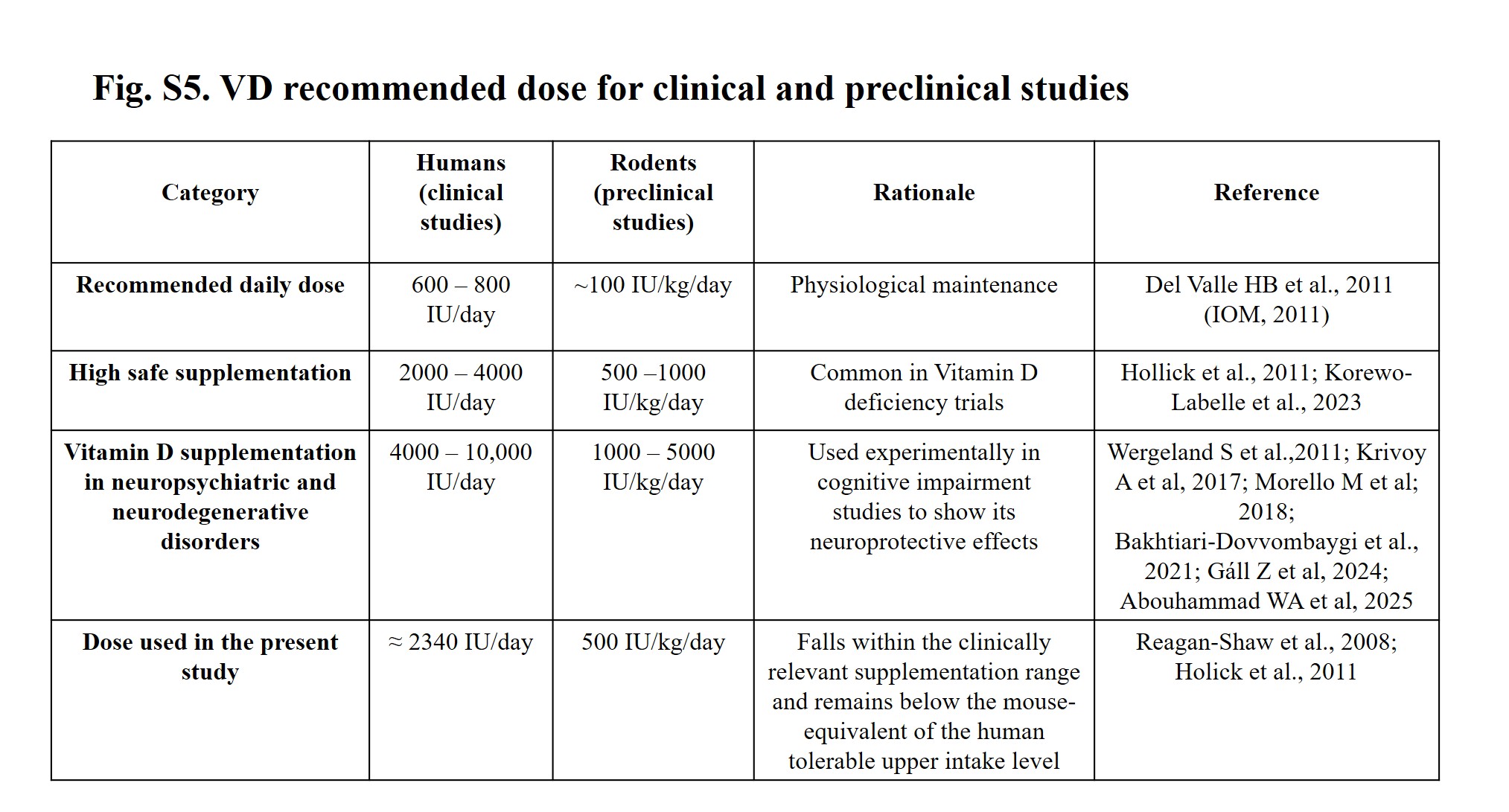
